# Metabolic constraints and quantitative design principles in gene expression during adaption of yeast to heat shock

**DOI:** 10.1101/143487

**Authors:** Tania Pereira, Ester Vilaprinyo, Gemma Belli, Enric Herrero, Baldiri Salvado, Albert Sorribas, Gisela Altés, Rui Alves

## Abstract

Microorganisms evolved adaptive responses that enable them to survive stressful challenges in ever changing environments by adjusting metabolism through the modulation of gene expression, protein levels and activity, and flow of metabolites. More frequent challenges allow natural selection ampler opportunities to select from a larger number of phenotypes that are compatible with survival. Understanding the causal relationships between physiological and metabolic requirements that are needed for cellular stress adaptation and gene expression changes that are used by organisms to achieve those requirements may have a significant impact in our ability to interpret and/or guide evolution.

Here, we study those causal relationships during heat shock adaptation in the yeast *Saccharomyces cerevisiae*. We do so by combining dozens of independent experiments measuring whole genome gene expression changes during stress response with a nonlinear simplified kinetic model of central metabolism.

This combination is used to create a quantitative, multidimensional, genotype-to-phenotype mapping of the metabolic and physiological requirements that enable cell survival to the feasible changes in gene expression that modulate metabolism to achieve those requirements. Our results clearly show that the feasible changes in gene expression that enable survival to heat shock are specific for this stress. In addition, they suggest that genetic programs for adaptive responses to desiccation/rehydration and to pH shifts might be selected by physiological requirements that are qualitatively similar, but quantitatively different to those for heat shock adaptation. In contrast, adaptive responses to other types of stress do not appear to be constrained by the same qualitative physiological requirements. Our model also explains at the mechanistic level how evolution might find different sets of changes in gene expression that lead to metabolic adaptations that are equivalent in meeting physiological requirements for survival. Finally, our results also suggest that physiological requirements for heat shock adaptation might be similar between unicellular ascomycetes that live in similar environments. Our analysis is likely to be scalable to other adaptive response and might inform efforts in developing biotechnological applications to manipulate cells for medical, biotechnological, or synthetic biology purposes.

## Introduction

Microorganisms evolved adaptive responses that enable them to survive stressful challenges in ever changing environments [1]–[8]. Adaptation to those challenges is achieved by adjusting metabolism to the new conditions, through the modulation of gene expression, protein levels and activity, and flow of metabolites [2], [8], [9]. Such adjustments integrate and balance the effects of stress with the physiological needs of the cell, ensuring that critical physiological parameters are tuned to guarantee survival [10]–[14]. One expects environmental challenges that were more frequently present during evolution to have selected for adaptive responses and metabolic adjustments of cells that are more finely tuned. More frequent challenges allow natural selection ampler opportunities to select from a larger number of phenotypes that are compatible with survival (***successful phenotypes***). Understanding this fine tuning and the qualitative and quantitative molecular determinants of stress responses may have a significant impact in our ability to interpret evolution, treat diseases, and manipulate microorganisms for medical, biotechnological, or synthetic biology purposes.

The yeast *Saccharomyces cerevisiae* is an important model organism for studying adaptive responses [15]–[23]. This yeast is, in many aspects, similar to more complex eukaryotes at the molecular level [15]. In addition, its genome, proteome, and metabolome are well characterized in a variety of physiological situations and there are many tools and methods available for manipulating and measuring the molecular responses of its cells. For these reasons we focus on that organism for the research reported here.

It is firmly established that the sets of yeast genes whose expression is modulated during adaptive responses to different types of stress only partially overlap [24], [25]. For example, *TPS1* and *TPS2* code for proteins involved in the synthesis of trehalose and change their expression in response to various types of stress [26]–[28], while *MEC1* only changes its expression in response to DNA damage, but not to heat shock [27], [29]. In addition, the changes in expression for ubiquitous stress responsive genes quantitatively depend on the type and intensity of the stress challenge, as can be seen by comparing various published experiments [28], [30], [31]. These quantitative dependencies suggest the existence of specific ranges for those changes that lead to ***successful phenotypes***, enabling cell survival. If this is so, yeast can only properly adapt to the specific challenge if it changes the expression of its genes within the boundaries established by those “feasibility regions” in gene expression space [10]–[14]. Investigating if such feasibility regions for gene expression changes exist and how and why they came about could allow us to understand their causal relationship with the physiological and metabolic requirements that are needed for cellular adaptation and survival. That understanding would create genotype-to-phenotype mappings of stress adaptation at the molecular level [10]–[13], [32]–[39].

To explore these issues we set our sights on yeast heat shock adaptation [13], [32], [33]. This adaptation occurs at different levels. For example, cell cycle stops, a new gene expression program starts, and the cell uses its ribosomes to synthesize specific protective molecules, such as chaperones. In addition, the cellular metabolism needs to be reorganized in order to permit survival and accommodate for all the changes at the level of gene expression and protein synthesis. The gene expression and protein synthesis levels of the heat shock adaptive response have been more thoroughly characterized and investigated than the biochemical and metabolic level (see for example [40]–[43] for reviews). Yet, it is known that the global adaptive response requires that central metabolism meets a varying set of physiological requirements for production of energy (ATP), reducing equivalents (NAD(P)H), and protective metabolites (e.g. trehalose or glycerol) [2], [10], [13], [33], [44]. These demands entail phenotypic adjustments of the levels and activities of proteins and metabolites, which could in principle be estimated from the modulation of gene expression via mathematical models [10], [13], [45]–[47]. Those mathematical models are a representation of the genotype-to-phenotype mappings of the cellular adaptation at the molecular level. They can be combined with experimental measurement of gene expression and used to identify the quantitative range within which metabolism and physiology must move for the cell to survive a specific stress challenge [10], [13], [14], [33]. These ranges, once known, can be used, together with the mathematical model, to solve the inverse problem of identifying the feasible regions for adaptive changes in gene expression that allow cells to adapt and survive [10]–[13]. The feasible regions for changes in gene expression represent quantitative design principles for genetic programs that generate appropriate adaptive responses to the relevant environmental challenges.

There are several problems to solve regarding the implementation of an analysis that permits identifying feasibility regions for heat shock adaptation. First, one needs to identify the parts of metabolism that need to be considered. Second, one needs to identify the physiological variables that are more adequate for establishing the feasibility regions of adaptation. Third, one needs to establish how (much) those variables must change to ensure adaptation. These problems were tackled by us [2], [13] and others [ 10], [40] before, using Monte-Carlo like simulations and global optimization methods.

Here, we extend that work to establish a systematic methodology that identifies quantitative design principles underlying metabolic adaptation based on gene expression profiles and apply it to the analysis of heat shock response in *S. cerevisiae* as a proof of principle. We adapt a minimal model of yeast central metabolism [10]–[13], [33] and use it to estimate the effect of changing gene expression on the production of energy (ATP), reducing equivalents (NAD(P)H), and production of metabolites that protect and stabilize cellular proteins and membranes, among other metabolic variables. We use this model and nine independent datasets (Supplementary Table I) from GEO [48] to estimate the feasibility regions for changes in gene expression and the quantitative physiological requirements that functionally constrain those regions. The quantitative boundaries for the feasibility regions of physiological changes are obtained from mapping the changes in gene expression (which we take as a proxy of the yeast’s genotype) to the changes in metabolism (phenotype) using the mathematical model. We then validate these quantitative predictions in two ways. First, we measure changes in gene expression in new heat shock adaptation experiments and find that they are consistent with the predictions. Second, we compare yeast adaptive responses for various types of stress and find that the feasibility regions and physiological requirements we identify are specific for heat shock response. These comparisons also reveal that our minimal model can be used to identify physiological constraints and feasibility regions that are specific for adaptation to desiccation/rehydration. In contrast, our model cannot be used to identify constraints and feasibility regions that are specific to the other types of stress response we analyze, indicating that additional metabolic variables likely impose physiological constraints that are specific for these responses. We conclude by discussing how to extend our analysis to other stress responses.

## Results

### Physiological requirements for cellular adaptation of *Saccharomyces cerevisiae* to heat shock

*S. cerevisiae* copes with heat shock by mounting a transcriptional response that modulates and adapts its physiology to the temperature increase. Overall, the eleven variables in Table I define quantitative physiological criteria that have been used by various groups in previous works to study and identify design principles of yeast metabolism during adaptation to heat shock [2], [10], [13], [33], [45]. In very simple terms, heat shock response requires that yeast increases production of ATP and NADPH (represented in Table I by variables V1 and V3, respectively), to allow for increases in the ATPase activity of the cell and to improve its reducing power, as one of the consequences of heat shock is an increase in oxidative stress [27], [29], [49]–[51]. Additionally, yeast should be able to produce enough protective metabolites to stabilize its proteins and membranes, such as trehalose and glycerol (represented in Table I by variables V2 and V10, respectively) [13], [45], [49]–[51].

**Table I:** Physiological variables used to identify possible design principles in the adaptive responses of yeast to heat shock.

| Variable | Acronym | Rationale for using this variable ([2], [10], [13], [33], [45], [150] and references therein). | Basal condition | Quantitative boundaries for heat shock survival |
| --- | --- | --- | --- | --- |
| V1 | V<sub>ATP</sub> | Changes in gene expression need to accommodate an increase in the rate of ATP production | 60 | 123.349<V1<339.19 |
| V2 | V<sub>Tre</sub> | Changes in gene expression need to accommodate an increase in the rate of trehalose synthesis | 0.0012 | 0.0076<V2<0.094 |
| V3 | V<sub>NADPH</sub> | Changes in gene expression need to accommodate an increase in reducing equivalents. The flux of NADPH production is used as a proxy for this increase | 1.77 | 4.31<V3<11.21 |
| V4 | GLC | Changes in gene expression should allow cells to avoid needless increases in the concentration of intermediates, thus minimizing possible toxic effects and the taxing of the solvent capabilities of the cell | 0.035 | 0.0094<V4<0.080 |
| V5 | G6P | Increases in the production/uptake of Glucose-6P are needed for the upregulation of energy production | 1.01 | 2.48<V5<19.91 |
| V6 | F16P | Depletion of Fructose-1,6,BisP needs to be tightly regulated and minimized, as this metabolite is an important bifurcation point in glycolysis that provides flux for the production of glycerol | 9.10 | 0.111<V6<20.58 |
| V7 | PEP | Changes in gene expression should allow cells to avoid needless increases in the concentration of intermediates, thus minimizing possible toxic effects and the taxing of the solvent capabilities of the cell | 0.0094 | 0.00019<V7<0.014 |
| V8 | ATP | ATP concentration should increase to meet energy demands | 1.12 | 2.39<V8<6.73 |
| V9 | Cost | Adaptation should be economic. We use changes in gene expression as a proxy for this variable. GEP (Gene Expression Profiles that allow adaptation with minimal changes in gene expression should be favoured | 0 | 8.10<V9<14.09 |
| V10 | V<sub>Glyce</sub> | Glycerol has a protective role in heat shock adaptation, and its production should either increase or not decrease by much. | 1.93 | 0.18<V10<2.07 |
| V11 | ψ | Changes in the activity of the enzymes TPS and PFK should be co-ordinately balanced after heat shock, in order to appropriately regulate the branching point in the glycolytic flux that divides material between glycolysis and trehalose production. | 52.06 | 5.34<V11<34.48 |

An important fraction of the material for the production of ATP, NADPH, trehalose and glycerol comes from glycolysis [49]–[51]. Thus, the flux distribution through the branching points of the glycolytic pathway is an essential target for fine tuning during heat shock adaptation. In broad terms, the activity of enzymes that draw material towards trehalose synthesis should be coordinately tuned with the activity of enzymes that produce F16P (represented by variable V11 in Table 1), while excessive depletion of F16P (Fructose- 1,6-bisphosphate – variable V6 in Table I) should be avoided in order sustain appropriate production of glycerol and ATP [2], [10], [13], [33], [45].

This global physiological adjustement needs to be made while taxing the solvent capabilities of the cell as little as possible. This can be achieved by buffering changes in the concentrations of unimportant metabolic intermediates as much as possible, which simultaneously contributes to prevent possible toxic effects of those intermediates [13], [33]. These changes in concentration can be measured by variables V4-V8 in Table I.

In addition, the adaptive response should be balanced and as economic as possible in terms of overall metabolic cost. A proxy that allows for a rough estimation of the relative costs caused by changes in gene expression during the adaptive response is given by variable V9 in Table I [2], [10], [13].

By using an appropriate mathematical model of metabolism one can estimate how experimentally determined changes in gene expression during response to heat shock propagate and change the physiological variables identified in Table I.

We note that the eleven variables from Table I do not constitute a full description of the whole heat shock response. For example, they do not explicitly consider the changes in chaperone ATPase activity that is characteristic of stress responses. Instead, they consider that change in a proxy manner, by estimating how ATP production changes and how cell uses its metabolic resources (Variables V1 and V9 in Table I). However, we show below that those eleven variables are sufficient to identify unique and specific quantitative requirements imposed on yeast by adaptation to heat shock.

### Minimal model for determining the physiological effects of changes in gene expression with respect to basal metabolism

In order to evaluate how changes in gene expression affect the central metabolism of yeast and the physiological variables detailed in Table I (among which production of energy [ATP], reducing equivalents [NAD(P)H], and protective metabolites [trehalose and glycerol] are especially important), we adapt and use a well-established mathematical model [10], [13], [33]. The system we model is described in more detail in Supplementary Figure 1 and in the methods section of the Supplementary Text.

In summary, the model accounts for a simplified version of glycolysis, for the oxidative stage of the pentose phosphate pathways, and for production of glycerol, trehalose, and NADPH. It was extensively validated as a good way to estimate the steady state values of the eleven variables from Table I [10], [13], [33]. The various individual processes in the model are catalysed by different enzymes. Each enzyme is coded by a (set of) gene(s) and, within the same pathway and as an approximation, the changes in gene expression can be used as a proxy for the changes in protein activity [52]–[55], which are represented in Eqs 1-5 by variables S1-S7 (see methods and Supplementary Table I for details). The rational for the modeling simplifications is also provided in the methods section. (see also [13], [33]). The model is used to estimate the basal values for the eleven variables in Table I, thus characterizing the basal steady state of yeast.

### Feasibility space for physiological adaptation of yeast to heat shock

In order to characterize the boundaries within which the genes considered in the model change their expression under heat shock, we extensively searched GEO for experiments that exposed *S. cerevisiae* to heat shock and downloaded 36 gene expression databases containing 410 experiments (Supplementary Table II). Out of these we selected all micro- or macro-array datasets that measure changes in gene expression occurring when yeast is shifted from temperatures below 30°C to temperatures above 30°C (HS datasets GDS15 [17-37°C, 21-37°C, 25-37°C, 29-37°C], GDS16 [25-37°C], GDS36 [29-33°C], GSE38478 [22-37°C]). We also used publicly available gene expression databases for yeast heat shock experiments (yeasts shifted from 25°C to 37°C) that are not available in GEO [28], [31]. All datasets are referenced in Supplementary Table II. The transcriptional changes of all the genes coding for enzymes in the model (Supplementary Table I) are then extracted from the resulting datasets, as described in the methods section.

The gene expression changes for each of the heat shock databases were used independently to estimate the changes in protein activities as described in methods. Those changes in protein activity were plugged into the model and the corresponding metabolic state under those new activities was calculated independently for each of the HS datasets. This allowed us to assess the approximate quantitative boundaries between which each of the variables from Table I can change to enable heat shock adaptation and survival. The results are summarized in Figure 1A, where variables labeled in red increase their value towards the center of the plot, while variables labeled in blue. The figure helps identify a well-defined region, marked in grey, within which the physiological adaptation of yeast to heat shock occurs, according to the eleven variables being estimated from the experimental results.

**Figure 1.**
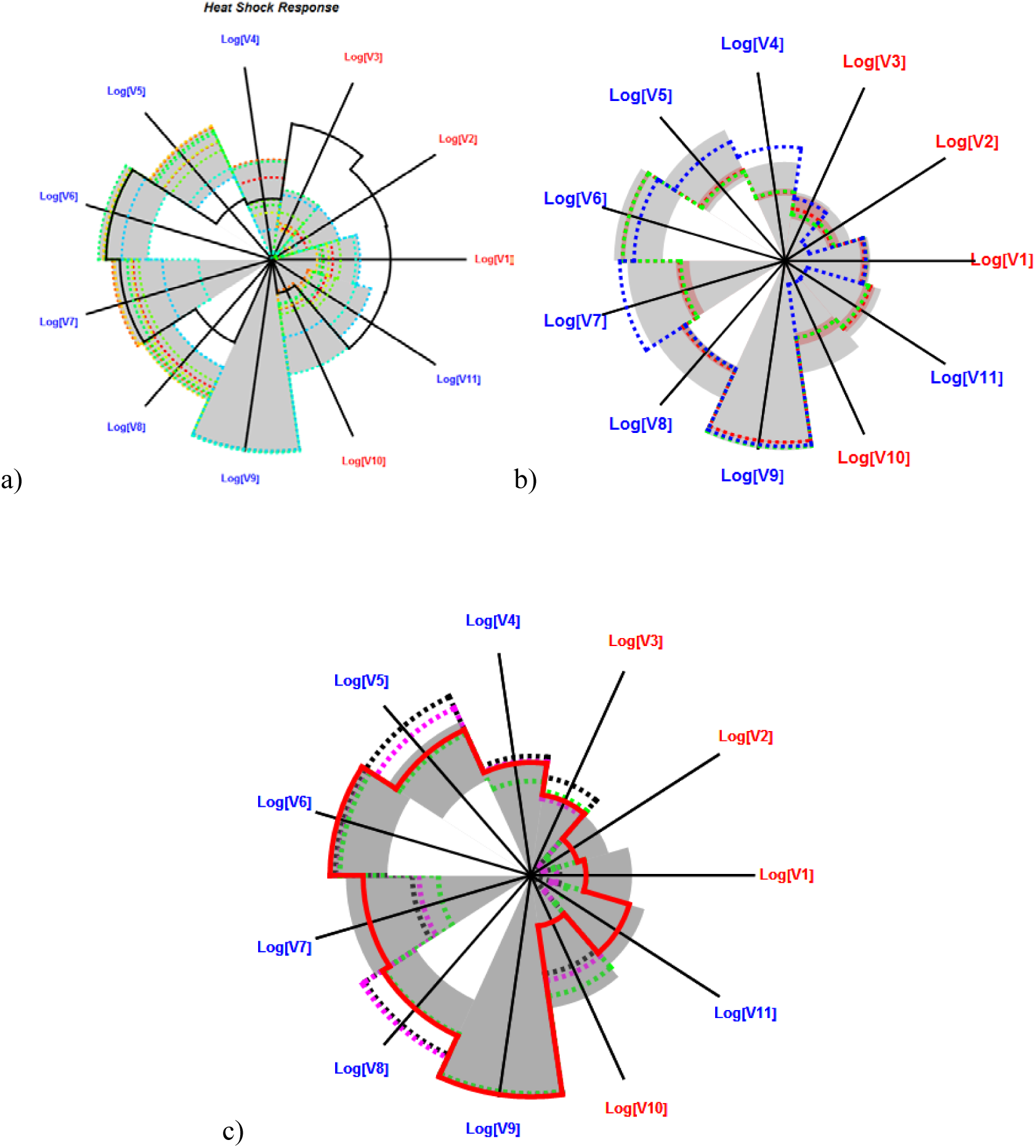
Spider plot representation of the feasibility range of adaptation of the eleven physiological variables from Table I during heat shock response. Each axis represents the logarithm of one of the variables. Variables in red grow towards the center of the axis. Variables in blue grow towards the outside of the axis. The grey area in all panels indicates the range of values that the eleven variables can assume in yeasts that adapt well to heat shock. The black line in all panels indicates the basal steady state values for each variable. A – Determination of the feasibility range using the heat shock experiments from supplementary Table II. Each dashed curve represents one of the databases; B – Validation of the feasibility range with independent experiments. The red and green curves represent the median and average (respectively) responses of our macro array experiment used to validate the feasibility range of the variables with a new yeast strain. The red area represents Quantiles 0.25 to 0.75 around the median determined using bootstrap. The blue line represents the values for the RefSeq experiment GSE58319 used to validate the feasibility range of the variables for a newer, more accurate, measurement technique; C – The red line represents the values for the eleven variables from Table I in response to a temperature shift from 29 to 33°C (GDS36). The dashed lines represent the values for the eleven variables from Table I for preadapted yeast that are subjected to a stronger heat shock (GDS15 – 33 to 37°C black line, GDS112 – 30 to 37°C magenta line, GDS2910 – 30 to 37°C green line).

Production of ATP (V1, see also V8, which is ATP concentration), trehalose (V2), and reducing equivalents (V3) is significantly increased during heat shock adaptation. This is easily seen because the basal values for these variables, which are represented by the full black line in Figure 1A, are smaller than those estimated during adaptation. In contrast, production of glycerol (V10) tends to decrease in our model, as indicated by the fact that the black line crossing the axis for variable V10 is below most of the other lines in Figure 1A.

Production of G6P (gucose-6-phosphate, V5) should increase to fuel increases in V1-V3 and V10. Figure 1A and Table I show that, in all cases, G6P is higher during heat shock response than at the basal steady state by about one order of magnitude. In turn, the concentration of glucose (V4) and PEP (V7) should be as buffered as possible against increases. In most cases, V4 decreases its value with respect to the basal steady state. When it increases, at most it doubles its basal value. In addition, V7 either remains close to its basal value or it decreases by at least an order of magnitude with respect to it. F16P levels (V6) are relatively buffered about the basal steady state value, changing by about a factor of two with respect to the basal situation. In addition, the changes in the variable that proxies for the cost of gene expression (V9) and for flux distribution in glycolysis (V11) also remain within a well-defined range.

Taking all the results together generates the grey region in Figure 1A. This region can be used as a proxy for the feasibility space of phenotypical adaptation of yeast to heat shock. We note that the smaller the fraction of the axis within the grey region, the smaller the range within which the corresponding variable must fall to ensure adaptation.

To ensure that the models for the various stress response databases were reasonable, we performed sensitivity and stability analyses on all calculated steady states. The sensitivity analysis showed that sensitivities to parameters and independent variables are small (see the results section of the Supplementary Text and Supplementary Table III), which implies that the computed steady states are quite robust to noise in the enzyme activities. This is to be expected for reasonable models of biochemical phenomena [56] and indicates that the models are robust to minor parameter changes. In addition, all steady states are stable, which is often a necessary condition for them to be physiologically relevant [56]. Details are given in the results section of the Supplementary Text.

### Validating the feasibility space for physiological adaptation of yeast to heat shock

The quantitative feasibility space identified in Figure 1A could be dependent on biological-environmental factors and on measurement techniques. To validate that space by changing a biological-environmental factor we performed additional heat shock experiments with a strain of *S. cerevisiae* that had not been used in the experiments we analyzed before to identify the feasibility space of Figure 1A. The new experiments are described in the methods section. We measured the change in whole genome gene expression for the yeast. The results for these experiments are summarized in Figure 1B. They are consistent with those obtained for the GEO databases, falling within the feasibility region defined in Figure 1A.

To validate the feasibility region by changing techniques, we searched for RNA-Seq experiments in GEO that also analyzed whole genome changes in gene expression during heat shock adaptation. Such experiments are reported in GEO datasets GSE58319 [57]. These datasets measure gene expression at mid-log growth phase, under basal conditions and after heat shock. According to our model, the changes in gene expression for these experiments lead to changes in variables V1-V3, V5-V6 and V8-V11 that fall within the feasibility region identified using array techniques (Figure 1C).Only variables V4 (glucose concentration) and V7 (phosphoenolpyruvate concentration) fall slightly outside of their feasibility ranges. These results suggest that feasibility regions might be a fundamental feature of adaptive responses that is robust to the measurement technique.

### Is the feasibility space for physiological adaptation to heat shock valid for pre-adapted yeast cells?

Yeasts can preadapt to a given stress by being exposed to small doses of that stress. The preadaptation process is also called hormesis [58]. Typically, pre-adapted cells are able to survive stress intensities that kill naïve cells [59]. The reason for the increased resistance lies on the fact that several protective metabolic adaptations are already in place and functioning when the new stress hits the cell, thus decreasing deleterious initial accumulation of cellular damage [13], [45], [59].

Taking this into account one might hypothesize that cells pre-adapted with a mild heat shock and then subjected to further temperature increases should, overall, still adapt their metabolism to fall within the feasibility space defined in Figure 1A for the eleven physiological variables.

To test this, we performed a two-step computational experiment. In the first step, we calculated how the values for the eleven variables changed in yeasts subjected to a mild heat shock. This was done by using experiment GDS36 [29 - 33°C] from Supplementary Table II to calculate the steady state of the cells in pre-adapted heat shock. The values for the eleven variables of this ***pre-adapted steady state*** are shown in Figure 1C. Ideally, a dataset for an experiment with the same strain and under the same conditions, but now subjecting the preadapted cells to an additional heat shock would be needed for the second stage of this experiment.

Given that no such dataset was available, and as an approximation, we selected three experiments from GEO where yeasts preadapted to between 30°C and 33°C were subjected to a stronger temperature increase (GDS15 – 33 to 37°C, GDS112 – 30 to 37°C, GDS2910 – 30 to 37°C). The second step of the experiment took the ***preadapted steady state*** calculated for experiment GDS36 as reference. Then, the mathematical model was used to estimate the variations in the eleven physiological variables of Table I caused by the gene expression changes reported in databases GDS15, GDS112, and GDS2910 (Figure 1C).

We see that most of the eleven variables fall within the feasibility regions identified in Figure 1A for the three experiments. The GDS15 experiment subjects cells to the mildest heat shock (4°C temperature shift vs. 7°C for the other two experiments). One would expect that the changes in the physiological variables for these cells should in principle be smaller than those observed for the other experiments. This would mean that the changes in variables V1-V11 should be closer to (or slightly on the outside of) the feasibility space boundaries in Figure 1C for experiment GDS15 than for experiments GDS112 and GDS2910. This is observed for all but two variables when we look at Figure 1C.

Experiments GDS112 and GDS2910 subject cells to a heat shock that shift temperature by 7°C and measure the time course of the changes in gene expression during adaptation. Thus, we expected that the estimated changes in variables V1-V11 are similar between the two experiments, which can also be confirmed in Figure 1C. We note that experiment GDS2910 has more replicates than GDS112, allowing for more robust estimation of changes in gene expression. This, in turn, made us hypothesize that the eleven physiological variables estimated for the GDS2910 dataset would be more robust and thus more likely to fully fall within the feasibility space estimated in Figure 1A.

The results are consistent with our predictions and hypothesis. All the eleven physiological variables calculated for the GDS15 experiment are either closer to and within the feasibility border (seven variables) or on the outside of that border (four variables). Only two variables calculated for experiment GDS112 are on the outside of the feasibility border and all variables calculated for experiment GDS2910 fall within the feasibility region of Figure 1A. Even with the approximations used in our two step computational experiment, the results are quantitatively consistent with the feasibility space determined in Figure 1A.

### Is the feasibility space specific for physiological adaptation to heat shock?

We wanted to understand how specific to heat shock is the feasibility space identified by our analysis in Figure 1A. In other words, are the boundaries for that space specific and valid only for heat shock adaptation or are they also applicable to adaptive responses to other stresses?

To answer this question we extensively searched GEO for experiments that exposed *S. cerevisiae* to various types of stress and downloaded gene expression databases related to such experiments (Supplementary Table II). These datasets measured changes in whole genome gene expression during yeast adaptation to desiccation, rehydration, osmotic, oxidative, reductive, and nutrient stress challenges. The transcriptional changes of genes coding for enzymes in the model were extracted from each dataset as described in the methods section, and the mathematical model was used to calculate how each of the independent sets of transcriptional changes affects the eleven constraints. Figure 2 summarizes the results. Specifically, Figure 2A shows that only heat shock responses fall within the feasibility region for all eleven variables from Table I. Figures 2B-2F also show that none of the curves that represent the changes in gene expression during the adaptive responses to other stress conditions fall fully within the feasibility space defined in Figure 1A. These results suggest that the feasibility space of adaptation shown in Figure 1A is specific for heat shock. They also suggest that variables V1-V3 are important in separating the adaptive response of yeast to heat shock from other adaptive responses.

**Figure 2.**
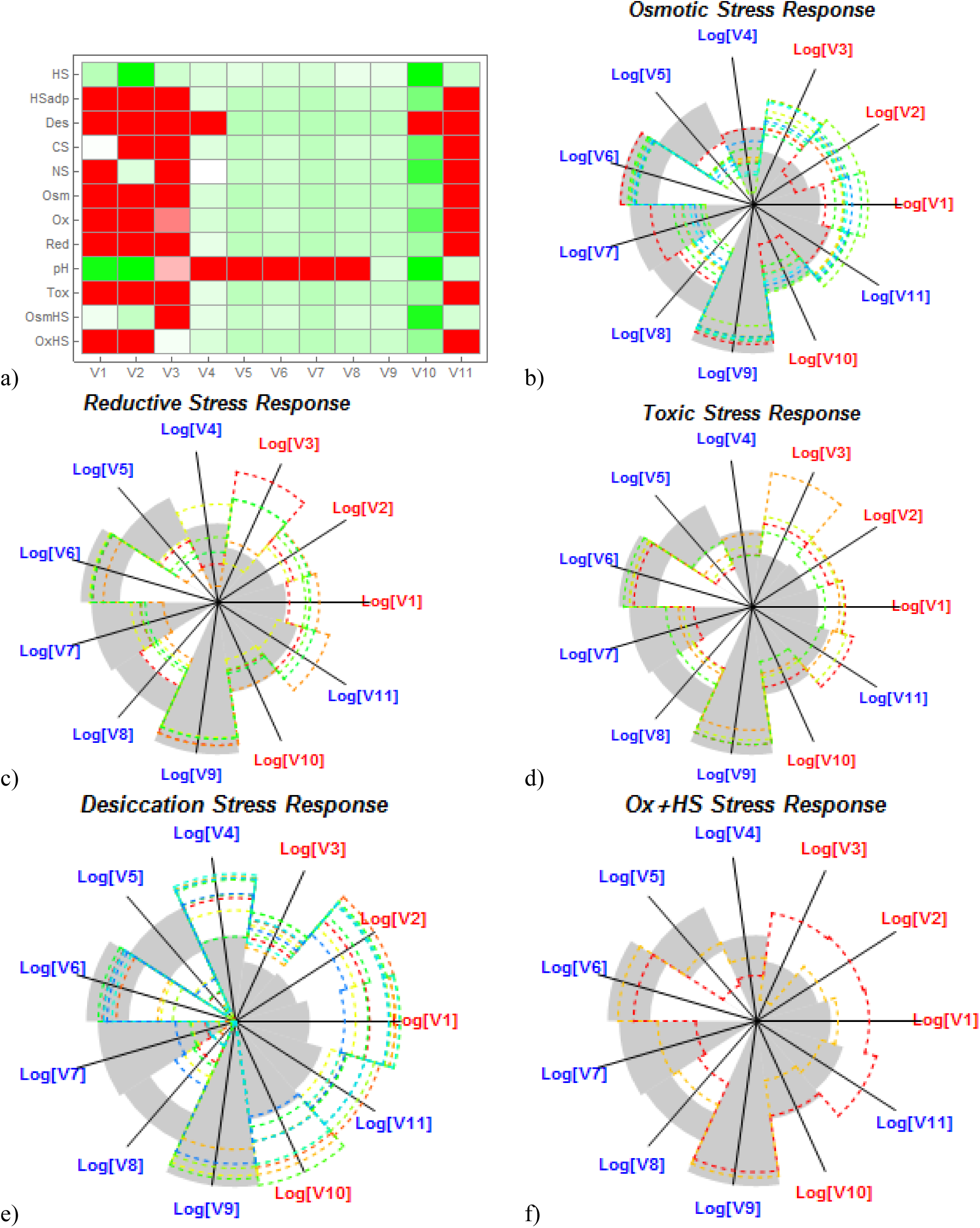
Specificity of feasibility space for heat shock adaptation. A – Summary of the results. Each row represents a type of stress and each column represents one of the physiological variables. Green (Red) entries indicate that the value of the variable falls within (outside of) the feasibility range for heat shock adaptation. More intense colors are further away from the feasibility boundaries. White indicates that the criteria are about the boundary value. Stresses: HS-Heat shock; CS-Cold shock; OxS-Oxidative; RS-Reductive; Osm-Osmotic; NS-Nutrient, Tox-Toxic, pH-pH stress, Des-Desiccation, SP-stationary phase. B – F: Spider plot representation of the adaptation of the eleven physiological variables from Table I. Each axis represents the logarithm of one of the variables. Variables in red grow towards the center of the axis. Variables in blue grow towards the outside of the axis. The grey area in all panels indicates the range of values that the eleven variables can assume in yeasts that adapt well to heat shock. Each curve represents an independent database from Supplementary Table II. B – Osmotic stress. C – Reductive stress. D – Toxic stress. E – Desiccation. F – Oxidative stress followed by heat shock.

### Can the same physiological variables be used to define specific feasibility spaces for other adaptive responses?

Our results indicate that the quantitative boundaries for the feasibility space are specific for adaptation to heat shock. Nevertheless, there is the possibility that the same physiological variables from Table I might be used to identify quantitatively different feasibility spaces for the adaptive responses to other types of stress.

To investigate this possibility, for each type of stress, we established the quantitative boundaries of the physiological changes observed for the eleven physiological variables defined in Table I, in the same way as we did for heat shock adaptive responses. This revealed that those variables create specific quantitative feasibility spaces for the adaptive responses to desiccation, rehydration, and (to a lesser extent) pH stresses (Supplementary Figure 2). These profiles are quantitatively different from those for heat shock. All other types of stress lead to feasibility spaces that are not specific. Hence, the eleven physiological variables studied here appear to be appropriate to identify quantitative design principles that are specific to heat shock, desiccation/rehydration, and pH stresses. Other variables need to be identified and used in a modified mathematical model for the remaining types of stress.

### Importance of individual physiological requirements

We identified specific quantitative boundaries for the feasibility space of physiological changes that allow yeast cells to adapt to heat shock, rehydration, desiccation and pH. We now ask which variables from Table I more strongly contribute to create a “signature of change” that uniquely identifies each type of stress response. Answering this question could help to establish which physiological variables more strongly contribute to the selective pressure that shapes adaptive responses as a whole.

We answered the question using feature subset selection, which is the process of identifying and removing as much irrelevant and redundant information as possible (see methods for details). Three independent methods for feature subset selection (Booth, Relief, and Correlation-based; see methods) identified the requirement for controlling cost of changing gene expression (V9) as the main variable for separating the adaptive responses to the various stress types, followed by the production rate of reducing equivalents (V3), protective trehalose molecules (V2), and energy (V1 and V8).

### Dependency between physiological variables

We established that the eleven physiological variables from Table I can be used to identify quantitative boundaries for adaptation that are unique for heat shock, and to a lesser extent for desiccation/rehydration and pH. We further showed that five of the individual requirements more strongly contribute to separate the various types of stress response. This number is not surprising, as the eleven physiological requirements are calculated from a model of five differential equations. The eleven requirements we use to calculate the feasibility space of physiological adaptation are a non-linear combination of the five dependent variables of the model. It follows that the eleven variables are redundant and interdependent. We now ask how many unique physiological variables we would need to separate the various stress responses.

To answer this question we performed the following experiments. First, we created an eleven-dimensional vector, where each dimension contains the value for one of the eleven variables defined in Table I. Second, we calculated the value for each variable V1-V11 in each stress response experiment from Supplementary Table I. Third, PCA (principal component analysis) was used to cluster the matrix of vectors.

This analysis was done independently three times. First, we performed it considering all stresses (Analysis SR1). Second, we performed it considering heat shock adaptation responses alone (Analysis SR2). Third and final, we performed it considering all stresses that were not heat shock adaptation (Analysis SR3). Results are summarized in Figure 3 and Table II. We see that heat shock responses are separated from the other responses mostly in the PC1 (principal component 1) dimension, while the Rehydration/Desiccation responses are separated mostly in the PC3 dimension (Figure 3 and Table II). Three principal components explain almost 90% (88.39%) of variability in the data for the SR2 analysis. The first principal component (PCA1) accounts for 63.57% of the variation in the data, the second (PCA2) 17.51%, the PCA3 only represents 7.31% and PCA4 represents 5.93%. The contribution of each variable to each of the PCs is shown in Table II. Variables that are the main contributors to PC1 and PC2 are similar in Analyses SR1, SR2, and SR3. The rates of ATP, trehalose, and glycerol production, together with concentration of pathway intermediates have the higher loadings in the PC1. In contrast, rate of NADPH production and metabolic cost of the changes have the higher loadings in the PC2. An important loading factor for PC3 in SR1-SR3 comes from variable V11, which evaluates the equilibrium between the flux of material that is used for energy production and the flux of material that is used for production of protective metabolites. In the SR2 Analysis the major loading for PC3 comes from variables V6 and V7, while variable V9 has the major loading in the SR3 analysis.

**Figure 3.**
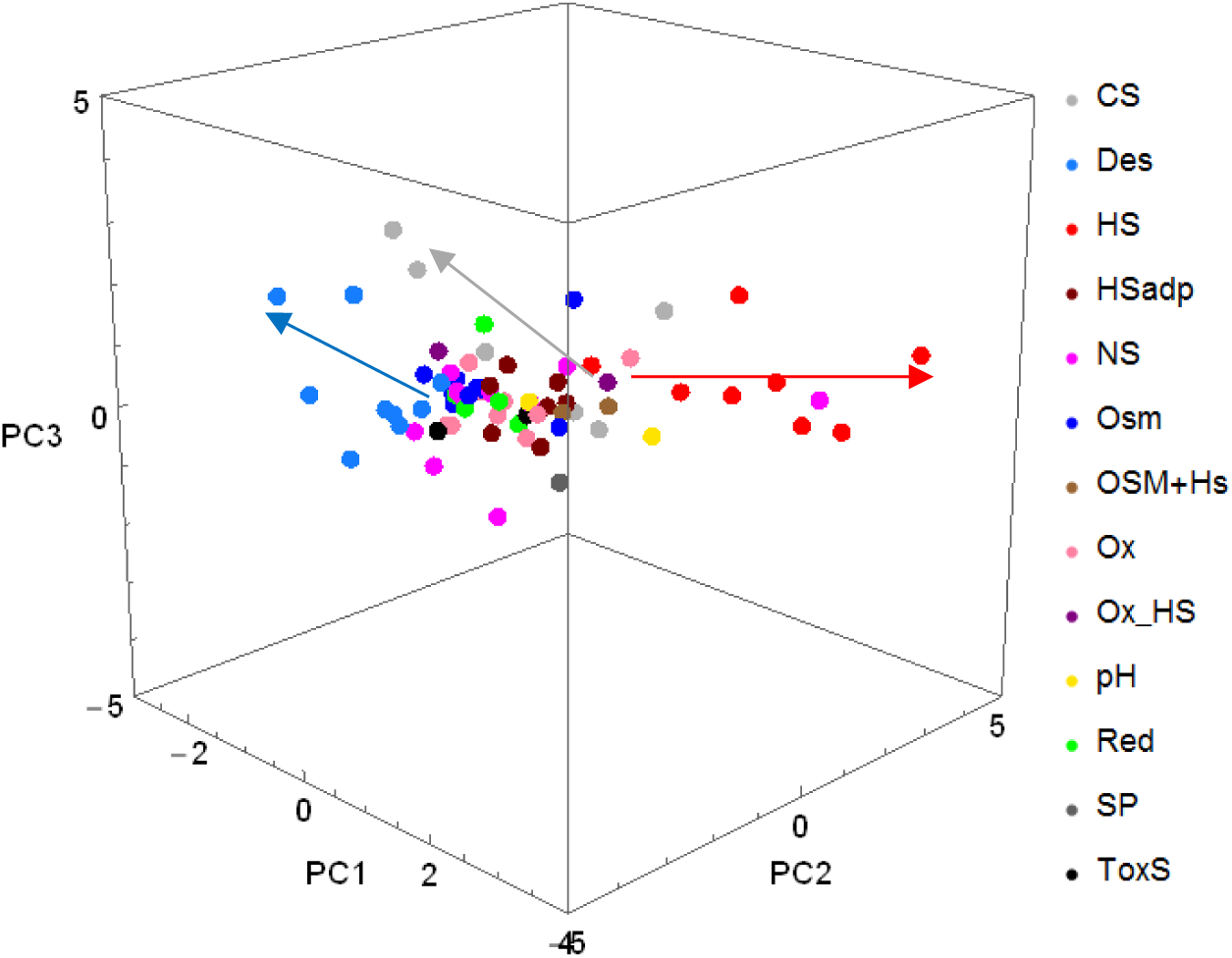
Principal Components Analysis of Variables V1 – V11 for all adaptive responses from Supplementary Table II. The top three principal components are represented. Each dot represents a sample, which is colored by stress type. PCA suggests that heat shock (red dots) and cold shock (grey dots) separate from each other and from other stresses mostly on PC1, while desiccation (blue dots) separates on PC3. Stresses: HS-Heat shock; CS-Cold shock; OxS-Oxidative; RS-Reductive; OsmOsmotic; NS-Nutrient, Tox-Toxic, pH-pH stress, Des-Desiccation, SP-stationary phase.

**Table II:**
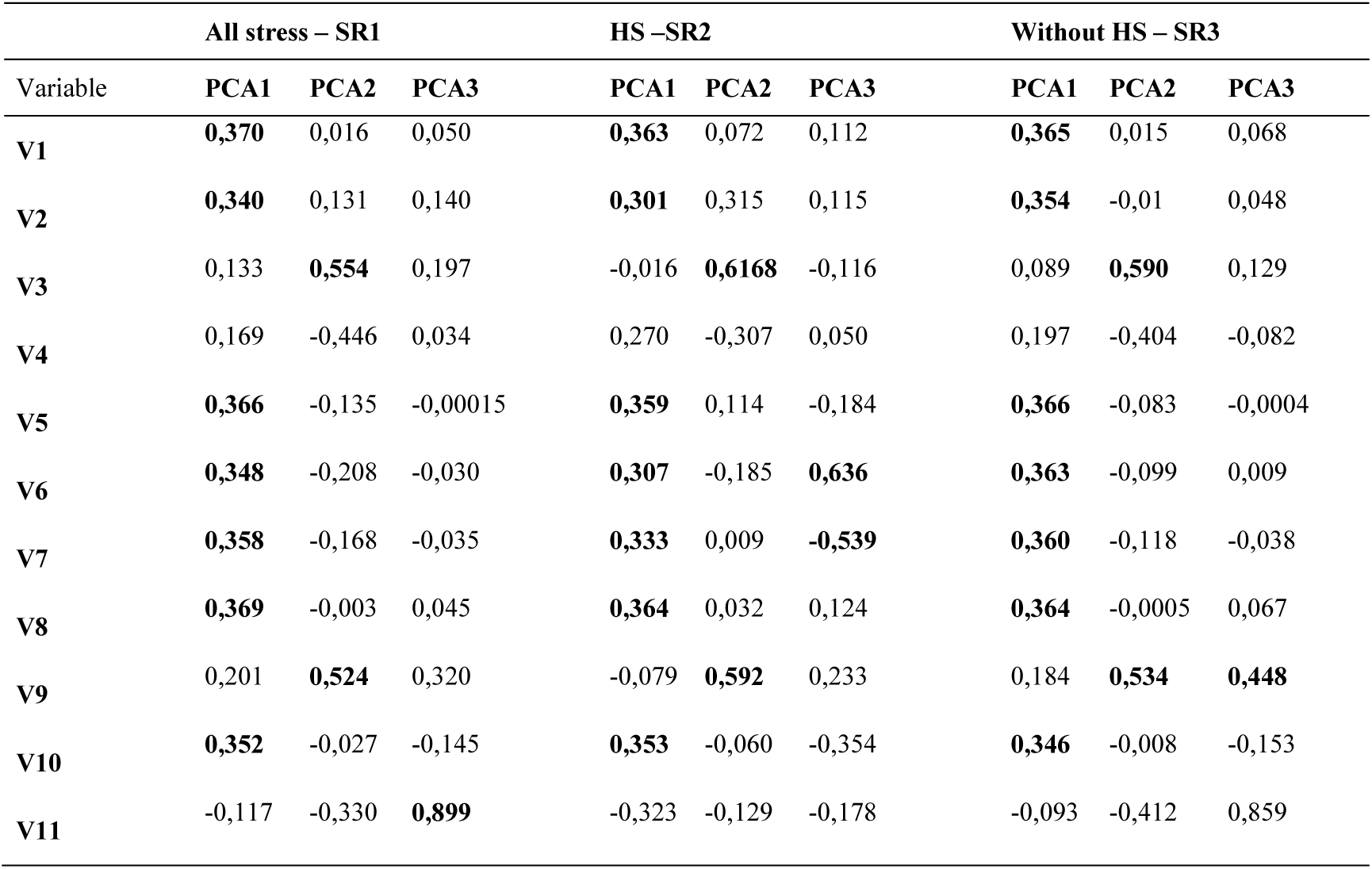
Variables used to identify design principles in metabolic adaptation of yeast to heat shock and their contribution to principal components.

### Mapping phenotype to genotype

The quantitative boundaries for the feasibility space of physiological changes was a result of mapping the changes in gene expression (genotype) to the changes in metabolism (phenotype) using the mathematical model defined in the methods section. This **Genotype-to-Phenotype** mapping is a one to one mapping. In other words, a set of changes in gene expression uniquely generates a set of changes in the physiological variables.

We used the same mathematical model to create an inverse mapping of the feasibility space for physiological changes to the corresponding feasibility space for changes in gene expression. This **Phenotype-to-Genotype** mapping is degenerate, in the sense that a set of changes in physiological variables can map to more than one set of changes in gene expression. Why is this so? The model considers seven independent enzyme activities (S1-S7) that are allowed to change in order to calculate the adapted physiological values of five dependent variables. When we solve the model with respect to those dependent variables, we obtain the steady state values for the adaptation, which is then used to calculate the eleven physiological variables from Table I.

If we now take the steady state for the dependent variables and solve the ODE system with respect to the enzyme activities, we can calculate five enzyme activities as functions of the dependent variables and of the remaining two enzyme activities. The calculations are shown in detail in the results section of the Supplementary Text, where we show that S1-S5 can be represented as linear functions of S6 and almost linear functions of S7. Given values for the activities of S6 and S7, and a range of values for X1-X5 one can then calculate the possible values for S1-S5. Figure 4A shows an example of the feasibility space of changes in enzyme activities S2 as a function of S6 and S7 for the extreme values of the dependent variables in the feasibility space of adaptation to heat shock (grey shade in Figure 1A). For the same steady state of the metabolic variables, the possible range of enzyme activities S1-S5 falls on a plane and depends on the exact value for S6 and S7 (see Supplementary Figure 3). Thus, cells can function at different values for the independent enzyme activities and still survive heat shock, if the differences between the activities of the various enzymes are coordinated in such a way that the values for the physiological variables that depend on those activities remain within the feasibility range for survival.

**Figure 4.**
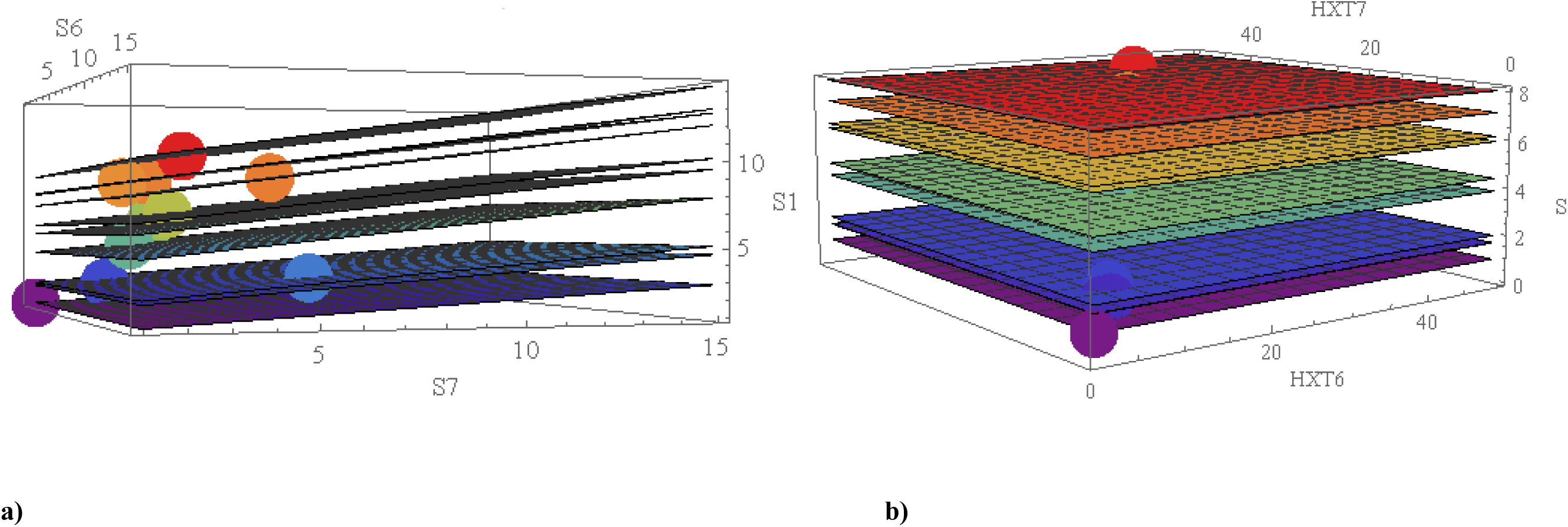
Evolution can find different combinations of changes in gene expression and enzyme activities that are equivalent with respect to the changes they cause in variables V1 – V11. Graphical representation of this situation for hexose transport activity S1. Each plane represents one of the heat shock response databases used to calculate the feasibility region shown in Figure 1A. A – Activity S1 as a function of activities S6 and S7. Each plane represents all possible sets of values for S1, S6, and S7 that would generate the same values for V1 – V11 for the same heat shock database. The dot that falls in each plane represents the actual activity estimated from the experimental changes in gene expression data. B – Activity S1 as a function of two high affinity transporters (HTX6 and HXT7) for the same heat shock responses we use as an example in panel A. Each plane corresponds to one of the databases. All points falling on a plane are formally equivalent, leading to the same S1 activity. The dots in each plane represent the actual measurement for the adaptive response.

A similar analysis can be done for the changes in gene expression. The enzyme activities represented by S1-S7 depend on a total of 22 genes. The mapping of the changes in enzyme activity to the changes in gene expression is defined in methods. S1-S7 depend linearly on the subset of the 22 genes that codes for proteins involved in the relevant enzyme activity. Figure 4B shows an example of the gene expression change to enzyme activity change mappings for the same examples represented in Figure 4A. Again, we can see that cells can use a wide range of coordinated changes in gene expression to adapt metabolism and make the value of physiological variables move to the feasibility region of adaptation. The representation for activities S1-S5 is shown in Supplementary Figure 3.

### How do heat shock responses of other fungal species conform to *S. cerevisiae*’s feasibility space?

The feasibility spaces and design principles identified so far result from evolutionary selection of metabolic adjustments, constrained by the physiological requirements that *S. cerevisiae* must meet to survive heat shock. Now we ask if the same (or similar) requirements can be extended to the adaptive responses of other unicellular yeasts to heat shock.

To answer the question we selected *Schizosaccharomyces pombe*, *Kluyveromyces lactis*, *Candida glabrata*, and *Candida albicans*. This selection was based on phylogenetic diversity and on availability of whole genome gene expression measurements during yeast adaptation to heat shock. The idea was to create species-specific mathematical models of the same pathways as those we model for *S. cerevisiae*. We would then use those models to estimate gene expression changes propagated to the eleven variables from Table I. After extensive literature and database searches, we could at best find kinetic information for only a small fraction of the reactions in the other yeasts. Therefore, it was impossible to proceed with the analysis in this way. Given that the few parameters we had found had the same order of magnitude as those in the *S. cerevisiae* model, we decided to use the *S. cerevisiae* model to perform the analysis.

The genes we considered in this comparative analysis were all those that code for protein in our mathematical model and have orthologs in the five yeast species (Supplementary Table IV). The GEO datasets from which we extracted gene expression data for each species are provided in Supplementary Table V [60], [61]. We used the same approach described in methods for *S. cerevisiae* to estimate how the measured changes in gene expression for the four non-*S. cerevisiae* yeasts would propagate to the enzyme activities. The estimated changes in enzyme activity were then plugged into the model for the basal state to calculate how the eleven physiological variables would change in S. *pombe*, *K. lactis*, *C. albicans* and *C. glabrata* (Figure 5). Estimated changes in the eleven variables in *S. pombe* and *C. albicans* fall within the feasibility space estimated for *S. cerevisiae*, with one or two minor exceptions.

**Figure 5.**
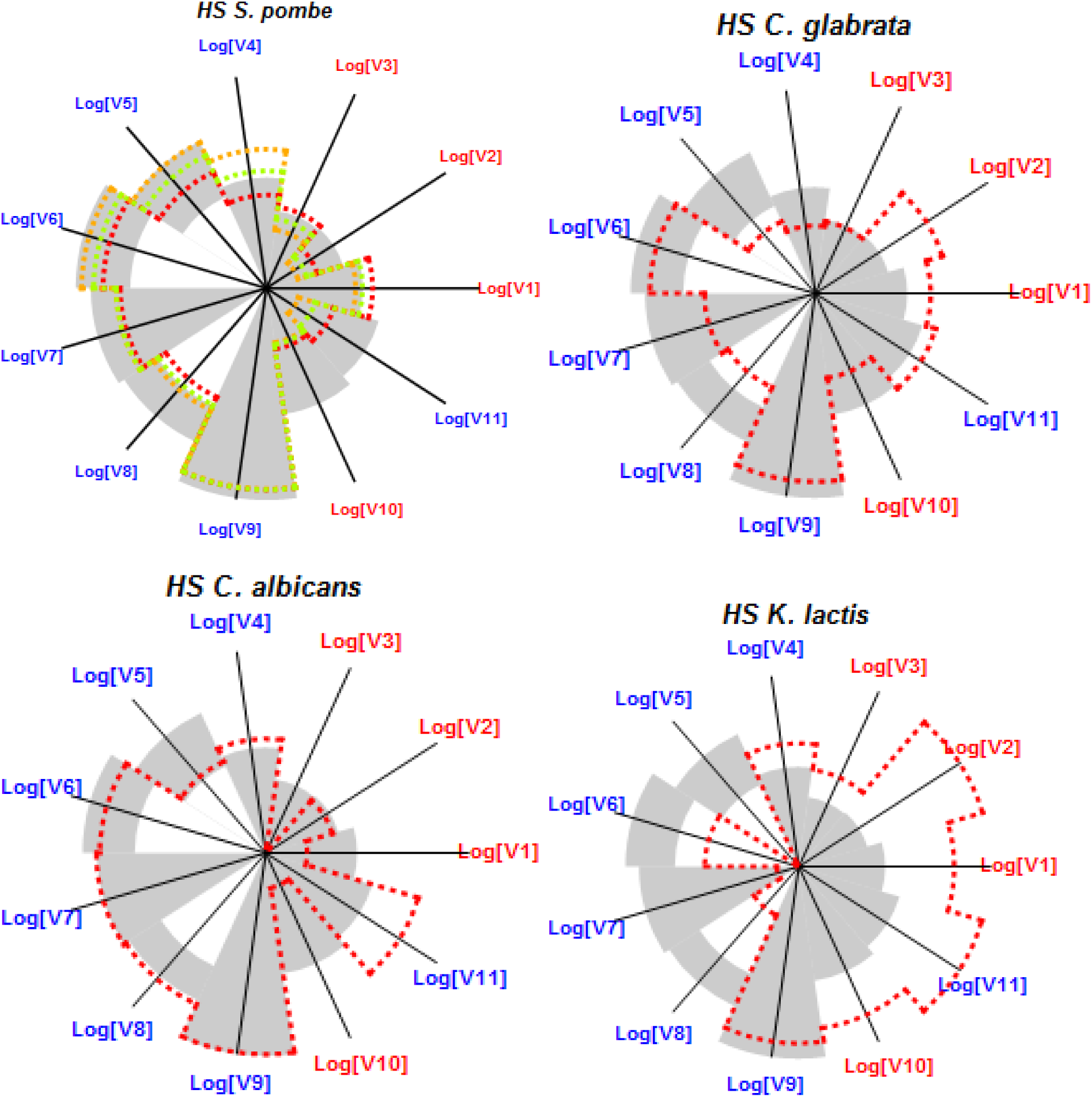
Spider plot representation of the feasibility range of adaptation of the eleven physiological variables from Table I during heat shock response for *S. pombe*, *K. lactis*, *C. albicans* and *C. glabrata*. Each axis represents a quantitative principle and the scale is logarithmic for all axes. Blue (red) axes indicate that the value increases as you move away from (towards) the centre of the plot. The grey area indicates the range of values for the criterion during heat shock in *S. cerevisiae*. A – *S. pombe*. B – *C. glabrata*. C – *C. albicans*. D – *K. lactis*.

The picture is somewhat similar for *C. glabrata*. While variables V1-V3 and V8 fall outside of the feasibility range for *S. cerevisiae*, they move towards that range and away from their basal values. This could have physiological justification. On the one hand *C. glabrata* is different from other yeasts in that it can only assimilate glucose and trehalose [62], [63]. This suggests that our *S. cerevisiae* model might not be a good estimator for trehalose turnover in *C. glabrata*. On the other, C. glabrata is a parasite that is accustomed to higher optimal growth temperatures than *S. cerevisiae*. This suggests that production of ATP and reducing equivalents might already be geared up for heat shock adaptation and that further adaptive changes to these variables could be smaller than for species with lower optimal growth temperatures.

When it comes to *K. lactis*, the picture is quite different. Our model estimates almost no change in the production of ATP, trehalose or reducing equivalents. This is not unexpected, as it is well known that *K. lactis* uses the mitochondrial tricarboxylic acid (TCA) cycle, respiration, and other non-conventional alternatives to produce reducing equivalents and ATP [64], [65]. When we analyse the changes in gene expression for respiratory complexes II-V in the microarray data, we find that between 20% (Complex III) and 35% (Complex IV) of the genes involved in each of the complexes are significantly upregulated in the microarray experiments. Furthermore, no gene involved in respiration is significantly downregulated. These results are consistent with an upregulation of respiration in *K. lactis* during heat shock adaptation [66]. Analysing the genes involved in the TCA, we also see that 20% of the genes involved in production of reducing equivalents are significantly upregulated, while no gene is significantly downregulated. Taken together, these results suggest a role of the mitochondria in production of energy and reducing equivalents. Given that our model does not account for mitochondrial processes, it is likely to be a poor estimator of the changes in the eleven physiological variables for *K. lactis*.

### Dynamical physiological adaptation of *S. cerevisiae* to heat shock

Most of the experiments we analyze compare the gene expression changes at an experimental end-point to the basal situation. Because of that, only a steady state analysis of the problem is possible. However, the transient dimension of adaptive responses is very important. In order to account for this and to understand how the feasibility space changes over time during *S. cerevisiae* adaptation to heat shock we focused on the nine databases from Supplementary Table II that contain time series information for gene expression changes (heat shock, nutrient starvation, osmotic stress, oxidative and reductive stress and desiccation). In addition, we use our own experimental datasets, which were generated to obtain more gene expression time series information (see methods).

In order to analyze how the eleven physiological variables change over time, and for each gene expression time series, we began by fully reconstituting the dynamics of gene expression using interpolation functions. Then, we used these interpolated time series as input for our model. This allowed us to simulate the transient shift of metabolism during stress adaptation (see methods for details). The results are summarized in Tables III and IV and given in detail in Supplementary Figure 4, where snapshots of metabolism are shown at 2, 10, 20 and 60 minutes after stress.

**Table III:** Ranges of dynamics values for each variable during adaptation to heat shock, as estimated by simulating the time course of adaptation. Initial values are those of the basal steady state.

| Time<br>(min) | V1 | V2 | V3 | V4 | V5 | V6 | V7 | V8 | V9 | V10 | V11 |
| --- | --- | --- | --- | --- | --- | --- | --- | --- | --- | --- | --- |
|  | V<sub>ATP</sub> | V<sub>Tre</sub> | V<sub>NADPH</sub> | GLC | G6P | F16P | PEP | ATP | Cost | V<sub>Glyce</sub> | ψ |
| 0 | 60 | 0.0012 | 1.77 | 0.035 | 1.01 | 9.10 | 0.0094 | 1.12 | 0 | 1.93 | 52.06 |
| 2 | 61.22-136.68 | 0.001-0.011 | 1.75-2.75 | 0.023-0.064 | 1.03-4.60 | 3.16-16.80 | 0.003-0.007 | 1.22-2.39 | 0.48-4.38 | 1.00-1.60 | 16.05-49.73 |
| 10 | 68.94-250.47 | 0.002-0.050 | 2.23-6.44 | 0.012-0.071 | 1.33-12.44 | 0.46-20.58 | 0.0009-0.0070 | 1.36-5.03 | 2.93-8.83 | 0.46-1.51 | 7.95-39.73 |
| 20 | 83.12-284.01 | 0.004-0.070 | 3.12-9.31 | 0.009-0.072 | 1.73-14.73 | 0.20-20.39 | 0.0005-0.007 | 1.61-5.73 | 5.10-10.72 | 0.32-1.49 | 6.68-34.50 |
| 60 | 98.61-309.11 | 0.005-0.090 | 4.31-11.16 | 0.0085-0.074 | 1.94-16.11 | 0.076-20.11 | 0.0002-0.007 | 1.70-6.26 | 7.02-13.23 | 0.17-1.49 | 6.00-36.36 |
| Ratio* | 1.39-4.73 | 3.33-58.99 | 1.76-5.26 | 0.26-2.06 | 1.71-14.58 | 0.02-2.24 | 0.05-0.74 | 1.44-5.12 | - | 0.17-0.77 | 0.13-0.66 |

\*Ratio between the ranges 20 minutes after the stress challenge and the basal value of the corresponding variable.

**Table IV:** How adaptive responses to various types of stress meet the feasibility space of heat shock adaptation for variables V1-V11. Each row represents one of the variables. Each column represents a time point in the dynamic adaptive response. Snapshots are taken at 2, 10, 20 and 60 min after stress. If a time point for a variable is left empty this indicates that all adaptive responses fall within the feasibility space defined by heat shock adaptation at that time (e. g. variable V1 at 2 min). Otherwise, the adaptive responses for the indicated stress at that time point falls outside of the feasibility range for the variable. HSadp – Dynamic response to heat shock by yeasts pre-adapted to a milder heat shock. NS+HSadp – Dynamic response to nutrient stress by yeasts preadapted to a mild heat shock. Ox – Dynamic response to oxidative stress. Red – Dynamic response to reductive stress. Des – Dynamic response to desiccation/rehydration. Osm – Dynamic response to osmotic shock.

|  | 2 min | 10 min | 20 min | 60 min |
| --- | --- | --- | --- | --- |
| V1 |  | HS<sub>adp</sub>, NS+HS<sub>adp</sub>, Ox and Des | HS<sub>adp</sub>, NS+HS<sub>adp</sub>, Ox and Des | HS<sub>adp</sub>, NS+HS<sub>adp</sub>, Ox, Red, Des |
| V2 | Red | NS+HS<sub>adp</sub>, Ox, Red and Des | HS<sub>adp</sub>, NS+HS<sub>adp</sub>, Ox, Red and Des | HS<sub>adp</sub>, NS+HS<sub>adp</sub>, Ox, Red, Des |
| V3 | Red | Ox, Red and Des | HS<sub>adp</sub>, NS+HS<sub>adp</sub>, Osm, Ox, Red and Des | HS<sub>adp</sub>, NS+HS<sub>adp</sub>, Ox, Osm, Red and Des |
| V4 |  |  |  | Des |
| V5 |  | HS<sub>adp</sub>, NS+HS<sub>adp</sub>, Ox, Red and Des | HS<sub>adp</sub>, NS+HS<sub>adp</sub>, Osm, Ox, Red and Des | HS<sub>adp</sub>, NS+HS<sub>adp</sub>, Ox, Osm, Red and Des |
| V6 |  |  |  |  |
| V7 |  |  |  |  |
| V8 | Des | HS<sub>adp</sub>, NS+HS<sub>adp</sub>, Ox and Des | HS<sub>adp</sub>, NS+HS<sub>adp</sub>, Osm, Ox, Red and Des | HS<sub>adp</sub>, NS+HS<sub>adp</sub>, Ox, Red and Des |
| V9 |  |  |  |  |
| V10 |  | Ox, Red and Des | Red and Des | Des |
| V11 |  | HS<sub>adp</sub>, Ox, Red and Des | HS<sub>adp</sub>, Osm, Ox, Red and Des | HS<sub>adp</sub>, Ox, Osm, Red and Des |

During heat shock response, production of energy (ATP) reducing equivalents (NADPH) and protective molecules (trehalose plus glycerol) sharply increases until twenty minutes after the heat shock, and tends to stabilize afterwards. Within the first twenty minutes the concentrations of ATP and G6P increase by a factor of two and four, stabilizing afterwards. The remaining metabolites remain roughly constant during the whole time course. The amount of resources invested by the cell in adapting metabolism to the new situation, as measured by variable V9, increases sharply for the first ten minutes of the response, remaining approximately constant afterwards. Similarly, at ten minutes, the cell reaches a new balance for resources allocated to the various synthetic branches of the model, as measured by variable V11. This global picture is reproduced when we use our own experimental measurements for gene expression changes in response to heat shock, which we also include in the time series analysis.

The timing at which the various variables from Table I reach the new steady state was similar for all types of stress we analyzed. However, the quantitative changes in energy production, NAD(P)H production, and trehalose production are always different from those observed during heat shock. In addition, the way that the glycolytic material is distributed between production of glycerol and ATP is also different between heat shock and the other stresses, as can be seen by comparing variables V10 and V11 in Supplementary Figure 4.

## Discussion

### Biological design principles

Biological circuits, pathways, or networks are conceptual models one uses to facilitate the understanding of how cells achieve a specific metabolic or physiological goal at the molecular level. These models are tremendously helpful for explaining in an organized way how cells perform the multiple tasks that help them survive. That conceptual formulation also has its limits, as often there is crosstalk between what one considers to be modular circuits or pathways. For example, every intermediate of glycolysis can be used by the cell in other pathways and many signaling pathways might crosstalk with each other. Nevertheless, it has been shown that biological networks tend to evolve modules that execute each of its required tasks in a fairly independent way [67]–[72].

This quasi modularity of biological circuits provides an opportunity for evolution to independently select and optimize each functional module that performs a specific task within the network. That selection may eventually lead to the spread in the population(s) of specific circuit designs that are the most effective ones in the environmental conditions under which they have evolved. In other words, they represent biological design principles that are optimal for the function they perform [73]– [80].

Many biological design principles for the structure of small biological pathways and circuits have been identified and explained in metabolic pathways, gene circuits and signaling pathways. For example, linear biosynthetic pathways are more efficiently regulated if the first enzyme of the pathway is inhibited by the final product of the pathway. This type of regulation provides for the best coupling between cellular demand for the product and flux through the pathway, the fastest response to changes in that demand and the least sensitivity to spurious fluctuations in the cell [81], [82]. In gene circuits, the demand theory for gene expression correlates mode of regulation for the expression of the gene with the fraction of the life cycle of the individual in which the gene product is needed [83]–[89]. Finally, in signal transduction, the design of the regulatory structure of bacterial two component systems could be selected based on whether the circuit needs to respond in a graded or switch-like manner and integrate signals from several sources or not [90], [91].

It also became apparent that these “hardware” design principles could be modified as additional individual components were integrated in a circuit, which itself could be integrated in larger networks [92], [93]. In addition, “software” or operational design principles were also identified [10], [45]. These operational design principles refer to how the parameters (concentrations and/or activities) of the individual components in a network might best be modulated to adjust environmental shifts. To our knowledge, these operational design principles have first been investigated in stress responses [10], [13], [33], [94], [95]. That research identified the first examples of feasibility spaces for modulating parameters that ensures adaptation and survival of the cell to various environmental challenges [13], [14], [32], [33].

### Evolution of adaptive responses

Stressful environmental changes required cells to evolve adaptive responses that ensure an appropriate reallocation of cellular resources in order to deal with and survive the insult [96]. Over time, natural selection favored those responses that more adequately enabled that survival (for example, see [97]–[99]). At the mechanistic level these responses favor dynamic gene expression programs that, together with post transcriptional regulation of protein activity, ultimately permit metabolism to suitably reallocate cellular resources during the response [100]. Given that functional redundancy between genes is very frequent in the genomes of free living organisms [101]–[104], many different gene expression programs might produce equivalent phenotypes. For example, imagine that cells need to produce more of a given metabolite that can be synthesized by two alternative reactions to survive a change in osmotic pressure. One subpopulation of cells might increase that production by upregulating genes for one of the reactions, while another subpopulation could upregulate the genes for the other reaction. There is ample evidence for this phenomenon in long term evolutionary experiments using unicellular and multicellular organisms (see for example [98], [105]–[108]). Thus, it is not trivial to predict the changes in gene expression that more adequately permit cells to adapt.

The evolution of such gene expression programs that are specific for a given type of adaptive response is fundamentally constrained by basal metabolism. Natural selection acts on the biological variability observed in the gene expression programs that allow cell survival under basal conditions, whatever those conditions may be. Once that is taken into account, one is tempted to speculate that the evolution of well-defined and stress-specific gene expression programs is more likely to occur if the type of stress underlying the evolution of that response is strong and frequently “seen by the cell” during evolution. This view is consistent with the Demand Theory for gene expression [85], [109], [89]. That theory argues that the selection of regulatory modes of gene expression is more likely if the regulation is frequently necessary for the survival of the cell. In addition, one also expects that expression programs for adaptive responses to stress conditions that challenge different parts of metabolism might be mostly modular and only slightly overlap [68], [72], [110], [111].

### Quantitative adaptation of yeast to heat shock

In this work we focus on the adaptive response of the yeast *S. cerevisiae* to heat shock. Specifically, we study how the cellular biochemical phenotype and its gene expression program (genotype) map to each other during adaptation to heat shock. There is a multi-level molecular adaptation of yeast cells to temperature increases. At the genomic level, there is modulation of gene expression that induces the production of chaperones, heat shock proteins, metabolic enzymes, and antioxidant defense proteins [13], [112]. At the proteome level, the activity of pre-existing and newly made proteins is regulated, both by temperature and by other metabolic events associated with the temperature increase [10], [113]. Finally, at the metabolomic level, the production of small molecules and metabolites is adjusted in order to allow yeast to meet the new physiological demands imposed on the cell by the temperature increase [45], [114]–[116]. Taken together, these events protect proteins and cellular structures, enabling recovery of the cell after stress adaptation.

All these molecular changes impose physiological demands on metabolism that require adjusting energy production in order to account for the increased ATP consumption. In fact, it is well known that ATP production after heat shock is about five times greater than that of the basal steady state of the cell [117]. During heat shock adaptation yeast needs to produce more NADPH reducing equivalents, in order to deal with the biosynthesis and redox protection of cellular components [118]. This is consistent with the doubling of NADPH production rate that our model predicts, which was also found to occur experimentally [119]. Overall production of protective molecules such as trehalose and glycerol needs to increase in order to help stabilize membranes and proteins [117].

In addition, there are generic physiological demands that impose optimization criteria and constrain the ways in which cells can adapt and evolve. These criteria include keeping flux distribution balanced, not taxing the cell’s solvent capabilities [120], [121] and adapt its metabolism in ways that are as economic as possible [2].

Taking into account the various physiological demands imposed on yeast metabolism by heat shock, we defined the metabolic variables in Table I as possible descriptors that can be used to measure how yeast changes its metabolism (Table I) as it adapts to heat shock. In previous work [10], [13], [32], [33], some or all of these variables had been used to investigate the physiological boundaries within which the metabolism of *S. cerevisiae* had to work if cells were to survive heat shock. In the current study we generalize the previous analysis and establish the boundaries for the physiological variables using a mathematical model that integrates information from a large number of gene expression experiments done in independent labs. These boundaries are proxies of the quantitative constraints, or design principles, that heat shock adaptation imposes on yeast metabolism. We also used the model to identify the permissible ranges within which the cell can change its gene expression and protein activities and still adapt to heat shock.

Our analysis indicated that, globally, the feasibility regions for changes in gene expression and in physiological variables were specific for heat shock response. Gene expression databases for other types of stress led to metabolic and gene expression changes that fall outside of the feasibility regions for heat shock adaptation. In addition, our own microarray heat shock experiments generated gene expression profiles and metabolic changes in the yeast model that fall within the previously calculated feasibility regions for heat shock. Our results also suggest that the feasibility regions we identified are likely to be biologically meaningful and generalizable to other *S. cerevisiae* strains (we compared 11 different strains). Comparing the heat shock response between *S. cerevisiae*, *S. pombe*, *C. albicans*, and *C. glabrata* suggests that the feasibility regions might also be at least partially generalizable to other unicellular yeasts. The predicted changes in the variables for the other three species fall reasonably well within the feasibility space defined by our *S. cerevisiae* analysis, in spite of the diverse ecology and lifestyles of these yeasts. The exception is *K. lactis*. In this case, we believe that our model does not account for important contributions to the changes in the physiological variables, given that this is an exclusively respiratory yeast [64].

The eleven metabolic variables we defined can also be used to identify quantitative constraints that are specific for yeast adaptive response to desiccation /rehydration stresses. This is not surprising, as yeast must produce significant amounts of trehalose and glycerol to stabilize membranes and proteins when the water content of the cell changes [114], [122]. In addition, significant increases in production of energy and reducing equivalents are needed to support activation of membrane ATPases and protein chaperones to avoid denaturation of proteins [123]. In contrast, the same eleven variables can not be used to separate metabolic adaptation of yeast to the other types of stress being analysed. This is not entirely surprising, because it is likely that alternative metabolic variables might be more important in determining the quantitative constraints for metabolic adaptation to those stresses [2]. For example, this is consistent with a complete re-adaptation of metabolism that yeast must make to survive nutrient stress.

Having identified the quantitative design principles that must be met by the metabolic variables in Table I for appropriate yeast adaptation to heat shock, we worked on the inverse problem. In other words, we investigated how those quantitative constraints impose restrictions on the changes in gene expression that yeast can make in order to adapt to heat shock. In our case, this analysis could be done by solving the mathematical model with the enzyme activities as dependent variables and the metabolites as independent variables. These experiments emphasized that there are multiple possible solutions to this biological problem (Figure 4 and Supplementary Figure 3). Evolution can explore a multidimensional space of gene expression and find solutions that are equivalent with respect to the variables we consider here. This is again consistent with many experiments where organisms are subjected to selection for thousands of generations and reach a new stable genotype. Afterwards they are subject to selection to return to their original metabolic state only to find that the old metabolic state is reached via a new set of mutations that are different from the original wild type strains [98], [105]–[107].

Our analysis used a steady state approximation to estimate how the eleven physiological variables of interest change as yeast adapts to stress. This was mostly due to the fact that very few experimental time series were available for the changes in gene expression during stress adaptation. However, the dynamic aspect of adaptation is crucial [124] and changes in gene expression peak at about 10-20 min after yeast cells are shifted from 25°C to 37°C (heat shock conditions) [10], [13]. Because of this, we extended the analysis to the transient regime of the adaptive response whenever time series were available. In addition, we measured our own time series for the changes in gene expression in response to heat shock and used them in the analysis of the transient part of the adaptive response. This analysis emphasized that the bulk part of the adaptive response occurred at most 20 minutes after the heat shock, both at the genetic and biochemical level, which is consistent with decades of research on the subject [26], [43]. In fact, we find that rates of metabolic production mostly adjust within the first two minutes of the response, while the new homeostatic metabolite levels are reached at about 10-20 minutes after the stress stimulus. This is consistent with measurements that showed that glucose levels increased at about 10 minutes after the stimulus, while trehalose levels increased after glucose reached its new homeostasis [117].

We point out that the work presented here provides a proof of principle for a methodology to establish quantitative design principles for metabolic adaptation to stress. This methodology can be summarized as follows. First, the metabolites, fluxes, and other metabolic variables that are important for the response should be tentatively identified. Second, a model for the pathways that contribute to the changes in those variables is needed. Third, estimates of how the various activities in the model change in response to stress are required. Fourth, these estimates are used to predict how the metabolic variables change in response to stress and the metabolic changes are used to identify the feasibility space of the metabolic changes. This feasibility space for physiological adaptation can then be used, together with the model, to estimate the feasibility space for the changes in protein activity and in gene expression, thus allowing us to establish a multilevel (metabolic, proteomic, and genomic) set of feasibility spaces for adaptation to stress. To conclude this part of the discussion we note that our analysis could in principle be extended to include a complete and general non-linear kinetic model of metabolism that could be used to study all stress responses indistinctly. In fact, such an analysis was done using much simpler FBA (Flux Balance Analysis) models [125]–[127]. However these models limit the variables that can be analysed to steady state flux distributions, which strongly constrains the type of physiological variables that can be studied.

### Limitations of this study

There are several limitations to this study. First, we did not analyse the design principles for changes in levels and activity of chaperones activated during heat shock to protect yeast proteins from heat induced denaturation and aggregation. Adding these chaperones would have required a significantly more complex model, for which not enough quantitative data is available. Nevertheless, we note that these are indirectly considered when we estimate ATP production and consumption. If (When) the model is expanded to include chaperones, additional axes will be added to the spider plots from Figure 1 and new feasibility sub-regions will be defined for those chaperones.

Second, our model simplifies the various pathways that need to be considered in order to identify the quantitative boundaries for the feasibility regions of adaptation. In spite of this simplification, the model and the analysis appear to be precise enough to identify boundaries for the metabolic variables and gene expression changes that are specific for heat shock adaptation. We are currently expanding the model and including more details about the pentose phosphate pathway, the production of trehalose and the production of glycerol. In addition, to identify important pathways in an unbiased manner, we are using pattern recognition methods to identify which pathways are modulated as a whole in response to a given stress type. This analysis will point us to areas in metabolism that are likely to constrain the way in which the organism adapts to the relevant stress type, suggesting which pathways need to be included in a model of the relevant adaptive response.

Third, when extending the analysis to other yeasts we assume that the changes in the eleven physiological variables we analyse are driven by processes and kinetics that are approximately similar to those of *S. cerevisiae*. In order to be more accurate, we attempted to adapt the *S. cerevisiae* model for each of the four additional yeasts by searching the literature and modelling databases for parameter values, basal concentrations, and activities that were yeast-specific. However, we found partial kinetic information only for *S. pombe* and, to a much lower degree, *C. albicans*. The few parameters and concentrations we found were very close to those for *S. cerevisiae*. Because of that, we decided to maintain the original model and use it to estimate changes in the eleven physiological variables. The lack of kinetic information also prevented us from creating models that more accurately described trehalose turnover in *C. glabrata* and relevant mitochondrial processes in *K. lactis*.

Fourth, the gene expression data we use come from macroarray, microarray, and RefSeq experiments, rather than from more precise methods, such as q-PCR. This choice was premeditated for two reasons. On the one hand, by selecting these types of experiments we had a larger number of datasets that were directly comparable and had a similar type of bias. This allowed us to confirm that the feasibility regions we analyse are specific for heat shock adaptation, and that additional metabolic requirements must be considered if we are to identify design principles for metabolic adaptation of *S. cerevisiae* to other types of stress. On the other, this type of experiment analyses whole genome gene expression changes, which will allow us to identify other pathways that might be important to include in developments of the model, as mentioned in the previous paragraph.

Fifth, our approach assumes that metabolism equilibrates itself on a time scale that is much faster than that of mRNA and protein synthesis. That is why we mostly focus on steady state analysis of the metabolic variable. Nevertheless, there is extensive evidence that this is so [128]–[131]. In order to control for any biases that this assumption might have introduced in the analysis, we also analysed the time course of the adaptive responses, whenever gene expression time series were available for doing so. The results of the time course analyses are fully consistent with those for the steady state analysis.

Sixth, we assume that changes in gene expression are mostly proportional to changes in enzyme activity. Although this is not always true, extensive experimental evidence suggests that the assumption is valid for the set of genes (metabolic enzymes) we consider in the model [132]. In addition, we downloaded whole proteome quantification data from PAXDB [133] and PRIDE [134] and compared the amount of the proteins from our model before and after heat shock. We found that the changes in protein amount between the two situations were strongly correlated with those seen in mRNA (See the results section of the Supplementary Text).

## Materials and Methods

### Mathematical Model

In order to understand how the eleven variables constrain changes in gene expression during heat shock response, we created a minimal mathematical model of the parts of metabolism that affect those variables. We used the GMA (Generalized Mass Action) mathematical formalism (see the Methods section of the Supplementary Text for details.). This model includes a simplified version of glycolysis that can be used to calculate how a specific change in a given gene will affect the physiological requirement that was identified in Table I. The mathematical model (Eqs 1-5) we use is given by the following five differential equations:

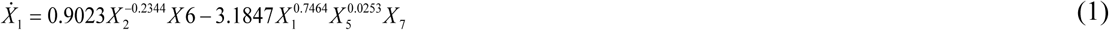

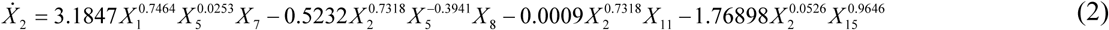

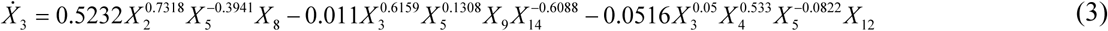

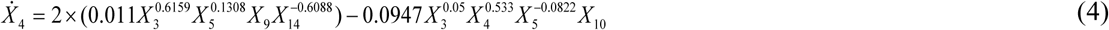

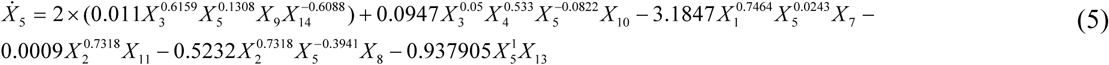

The dependent variables contained in the model are: X1 = internal glucose, X2 = G6P, X3 = F16P, X4 = PEP, X5 = ATP; and independent variables are: X6 = glucose transporter activity, 19.7 mM min-1; X7 = HXK/GLK activity, 68.5 mM min-1; X8 = PFK activity, 31.7 mM min-1; X9 = TDH activity, 49.9 mM min-1; X10 = PK activity, 3440 mM min-1; X11 = trehalose production activity, X12 = glycerol production activity, 203 mM min-1; X13 = ATPase, 25.1 mM min-1; X14 = NADH/NAD+ ratio, 0.042; X15 = G6PDH activity, 1. Parameter values and basal steady state concentrations are extracted from [33]. The model, its assumptions, and how parameter values were estimated is described in detail in [13], [33]. Calculating the changes in trehalose and NADPH production is done using the following equations:

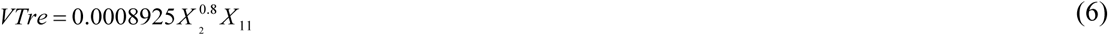

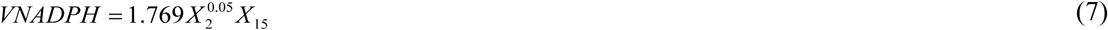

These equations account for the major known source of NADPH production in yeast during heat shock adaptation, which is the oxidative phase of pentose phosphate pathway [124]. Although Eqs. 6-7 are a simplification of more complex pathways, there is ample experimental evidence suggesting that these approximations are reliable [13], [33].

### Gene expression data

Experiments that exposed *S. cerevisiae* to stress and measured how the yeast adapts its gene expression were identified by first searching GEO [135] for “stress” and “cerevisiae” and then manually going through the list and identifying all experiments where classical stress challenges where given to any strain of *S. cerevisiae*. All such databases of micro array data were downloaded and stored locally. 38 databases, containing 81 different independent experiments were analysed (Supplementary Table II). Stress challenges included heat shock (HS), cold shock (CS), oxidative (Ox), reductive (Red), osmotic (Osm), desiccation (Des), pH, toxic elements (Tox), nutrient starvation (NS) and stationary phase (SP). All gene expression data were converted to the logarithmic ratio with respect to a basal pre-stress condition. Four additional databases were excluded, because only absolute determinations of post-stress mRNA abundance were given and no information regarding basal gene expression was found (GDS2969, GDS2050, GDS991, GDS777). Two other classical databases, not included in GEO, were added to this study [28], [136].

### Estimating changes in gene expression

All entries for the twenty two different genes considered in the model were extracted from each database. The gene expression databases used in this work resulted from three types of experiments: replicated measurements of an end-time for the adaptation, independent conditions of experiment (some databases are composed by experiments with different conditions and for this reason were analysed separately), and time course experiments. For each gene, and at each time point, we eliminated missing values and then used the average of the remaining entries as the representative measure for the change in gene expression. Additionally, for time course experiments, we needed to estimate whether a gene was globally activated or repressed, as a single value was required to calculate the steady state. This was done in the following way. The logarithmic changes in expression for the gene were plotted as a function of time. The area under the curve in the positive quadrant of expression was divided by the area under the curve in the negative quadrant of expression. If the ratio was larger (smaller) than one, we considered that the gene was overexpressed (repressed). The quantile 0.9 (0.1) of change in gene expression over the time course was calculated for overexpressed (repressed) genes, and this number was used to estimate the global change in gene activity. The quantiles were used instead of maximum or minimum in order to decrease the probability of using outlier values.

### Estimating changes in enzyme activity

All genes coding for proteins directly involved in the enzyme activities of the model were considered, These were: S1 – hexose transporters - HXT (HXT1, HXT2, HXT3, HXT4, HXT6, HXT8); S2 – glucokinase/hexokinase - GLK (genes: GLK1, HXK1, HXK2); S3 – phosphofructokinase - PFK (PFK1, PFK2); S4 – glyceraldehydes-3-phosphate dehydrogenase - TDH (TDH1, TDH2, TDH3); S4 – pyruvate kinase - PYK (PYK1, PYK2); S6 – trehalose synthase complex - TPS (TPS1, TPS2, TPS3) and S7 – glucose-6-phosphate dehydrogenase - GD6PDH [10].

We could not find direct measurements for the changes in all enzyme activities of the model under heat shock. Nevertheless, it is well documented that changes in enzyme activity and gene expression are highly correlated in glycolysis [132], Because of this we assumed that the fold-change in gene expression directly translates into a similar fold-change in the activity of the corresponding enzymes, whenever a single gene coded for that enzyme activity.

If more than one gene contributed to an enzyme activity we also assessed the relative contribution of that gene to the enzyme activity. In cases where the genes coded for alternative enzymes that have the same activity, we weighted the change in gene expression by the basal abundance of the protein using the formula:

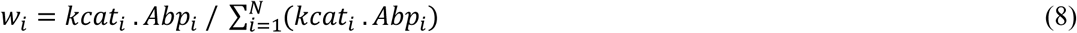

In Eq 8, Abpi represent the basal abundance of protein *i* in the cell and k_cat_ represents the catalytic constant of the enzyme (Supplementary Table I). We note that *kcat* for hexose transporters were not found in the literature. The relative contribution of the alternative transporters was estimated using the information of V_max_ and the concentration of molecules per cell [ 137].

The expressions used to calculate the new activities of S1 – S7 for each database are

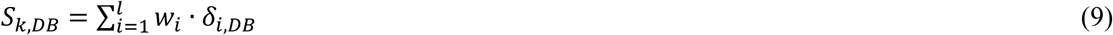

In Eq 9, *w_i_* are the weights for each gene, as defined by Eq 8, and *δ_i,DB_* represents the change in gene expression for the relevant gene in experiment DB. We note that the changes in PFK activity estimated using Eqs. 8-9 are further multiplied by a factor of five to account for a five-fold posttranscriptional activation of this enzyme during heat shock response [13]. Similarly changes in TDH activity estimated using Eqs. 8-9 are further multiplied by 1.5 to account for a 1.5-fold posttranscriptional activation that is also known to occur during stress response [13].

The information about the kinetic properties of the various enzymes was compiled and is presented in Supplementary Table I. We note that *in vivo* measurements for the parameters were used whenever available. When only *in vitro* determinations were found, they were used as a proxy of the *in vivo* values, because it was shown that both values correlate [138].

### Finding orthologs in other yeast species

Orthologs for the twenty two *S. cerevisiae* genes in *Schizosaccharomyces pombe, Kluyveromyces lactis, Candida glabrata and Candida albicans* were identified using Uniprot [139], searching for the species name combined with the words: heat shock. These orthologs are given in Supplementary Table IV.

### -Macroarray experiments

*S. cerevisiae* wild type strain W303-1A was employed for the determination of mRNA levels upon heat shock. Cells were grown exponentially in YPD medium at 25°C, at time 0 they were quickly shifted to 37°C by dilution with three volumes of pre-warmed fresh medium at 41°C, and then maintained in a 37°C water bath for subsequent recovery of samples at different time points. Four independent experiments were carried out, and for each experiment two samples were processed for each time point (eight replicates per time point). Total RNA isolation and labelling, and determination of mRNA levels were done as described in [23] at 0, 3, 6, 9, 12, 15, 18, 21, 25, 30, 45, and 60 min after heat shock. Values at each time point after the beginning of the experiment were normalized by those at time 0.

Bootstrapping was used to determine confidence intervals for the changes in gene expression at each time point in the following way. Four replicates were randomly selected from the eight experiments one hundred times. The average time series for each set of replicates was estimated. Then, we calculated quantiles 0.025 and 0.975 of the bootstrapped datasets to estimate the 95% confidence interval for the changes in gene expression at each time point.

### Steady state robustness

Biological systems must be able to adapt to and survive in an ever-changing environment, without being overly sensitive to small changes that are spurious. To achieve this, most biological systems have low sensitivity to fluctuation in parameters (e.g. enzyme activity or *Km*) and such fluctuation will not greatly affect its steady state or homeostasis [140]–[142]. This is called robustness of the steady state and it can be measured using sensitivity analysis [140]–[142]. In this work we evaluated the robustness of the physiological variables V1-V11 using relative steady state parameter sensitivities (see the Methods section of the Supplementary Text for details). In our case, we wanted to determine the relative sensitivity of the physiological variables V_1_ – V_11_ of Table I to the enzyme activities S1 – S7. The results of this sensitivity analysis identify the enzyme activities that more strongly influence each of the relevant physiological variables. In approximate terms, if 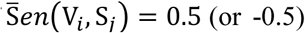, this means that when the value of S_j_ changes by 100%, the value of V_i_ is expected to increase (or decrease) by approximately 50%.

### Steady state stability

A biologically meaningful steady state is stable, and homeostasis should be quickly re-established if the system fluctuates away from it. For example, after finding a small bolus of nutrients, this bolus should be consumed and the original steady state is reinstated, if it is stable. This property of the steady state is called stability and we analysed the stability of each steady state using stability analysis (see the Methods section of the Supplementary Text for details).

### Principal component analysis

Principal component analysis (PCA) is a method to reduce the dimensionality of a dataset and identify which orthogonal linear combinations of variables contribute more strongly to the quantitative variation in the data. PCA of the matrix containing the eleven metabolic variables for each stress experiments was done in the following way: was determined the correlation matrix, from this matrix was calculated the *eigenvalues* and *eigenvectors*. The eigenvalues represent the amount of variation explained by each factor. A *varimax* was used as a redistribute process that can help interpret the principal components, now containing loadings towards either +1 or -1 [143].

### Feature Analysis

While PCA provides a way to determine how many dimensions one needs to describe the variability in the data at various degrees of accuracy, often the PC themselves are difficult to interpret. Because of that we also performed feature analysis. Feature analysis was done using two methods: Relief-based feature selection (RFS) and Correlation-based Feature Selection (CFS) RFS works by randomly sampling an instance from the data and then locating its nearest neighbour from the same and opposite class. The values of the attributes of the nearest neighbours are compared to the sampled instance and used to update relevance scores for each attribute [144]. CFS works by evaluating the subsets of attributes rather than individual attributes, and takes into account the usefulness of individual features for predicting the class along with the level of inter-correlation among them [145].

### Analysis of the Transient Response

The temporal dynamics of the model was studied in all cases where time series were available for the gene expression data (databases GDS16, GDS20, GDS30, GDS31, GDS34, GDS36, GDS108, GDS112, GDS113, GDS2712, GDS2713, GDS2715, GDS2910, GDS3030, GDS3035 and GSE38478 from Supplementary Table II). We simulated the adaptation of yeast to stresses from zero to ninety minutes after stress. This analysis was done in the following way. First, the time series for each gene were taken from time zero until the maximal change in gene expression was observed. This value was then maintained for the remaining of the time series. This modified time series was then represented using Akima interpolation, thus guaranteeing smooth and differential curves [46], [146]. This procedure aims at capturing the difference in time scales between changes in gene expression (faster) and changes in protein synthesis and degradation (slower) [27], [147]. The interpolated function for each gene was then used to calculate the changes in protein activity at each time point, as described in subsection “Estimating changes in enzyme activity”, above. These values were then fed into the differential equations and simulation was used to estimate the dynamics of the eleven variables in Table I.

### Software

Most calculations were done using Wolfram Mathematica [148]. Databases of gene expression were also imported and manipulated using Mathematica. The steady states for the system of ordinary differential equations were solved using the *FindRoot* function. Time course simulations were done using the *NDSolve* function [148]. Graphical representations were also done using plots designed in-house with Mathematica. PCA (Principal component analysis) was also done using Matlab™ and feature analysis was done using Weka 3 [149].

## Acknowledgements

This project has received funding from the European Union’s Seventh Framework Programme for research, technological development and demonstration under grant agreement no [609396]. This work was partially funded by grants BFU2008-0196 and BFU2010-17704 from the Spanish MINECO, 2009SGR809 from Generalitat de Catalunya, and bridge grants from the Dean for Research and the Departament de Ciències Mèdiques Bàsiques of the University of Lleida.

## Author contributions

EV, AS and RA conceived the project; GB and EH designed and performed the microarray experiments; TP, EV and RA performed the computational experiments. TP, EV, RA, BS, GA and AS analysed the results. All authors contribute to write various iterations of the paper and approved the final manuscript.

## Supplementary Figure Captions

**Supplementary Figure 1.**
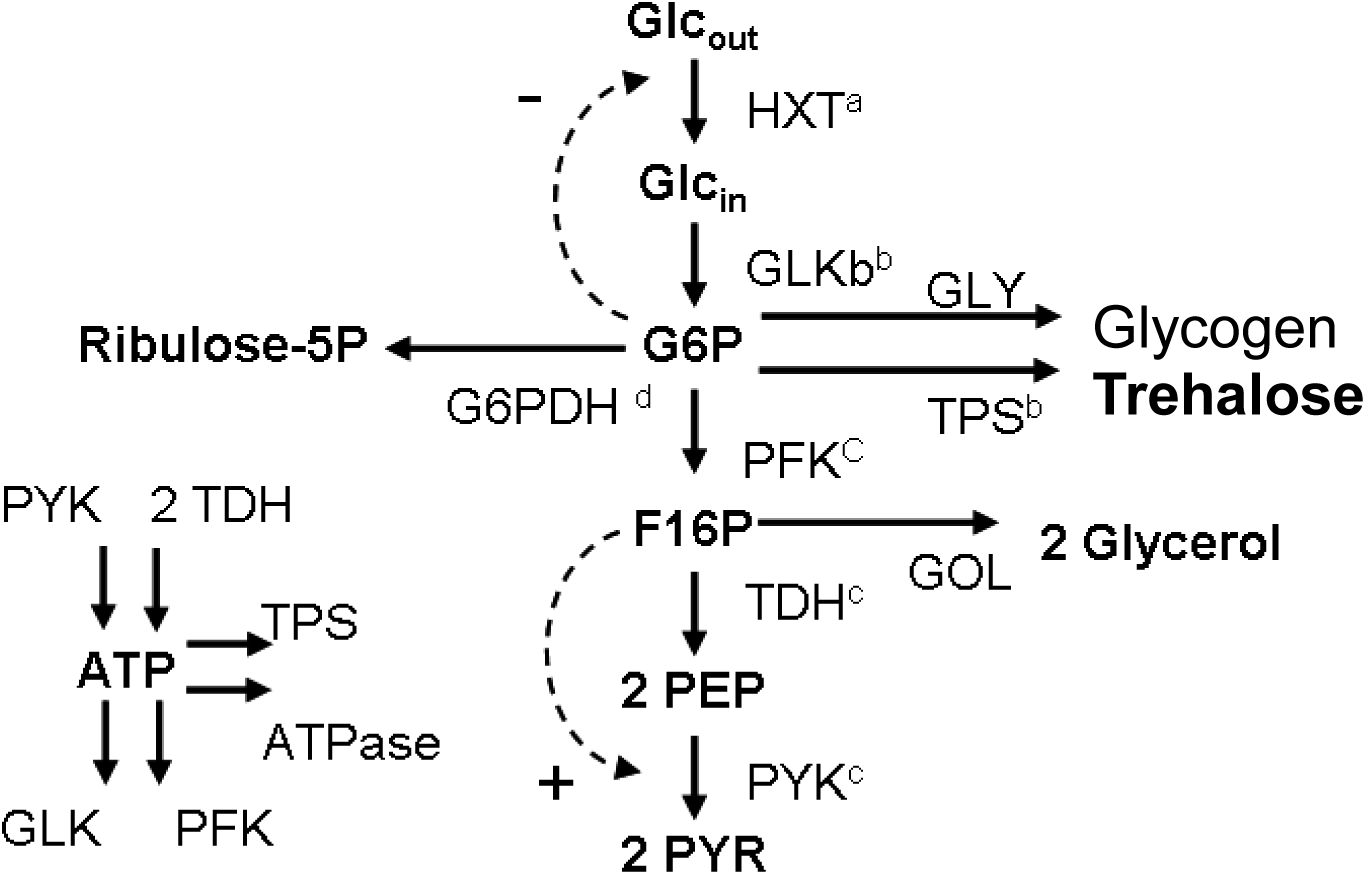
Minimal model to account for the metabolic changes described in Table 1 of the main manuscript.

**Supplementary Figure 2.**
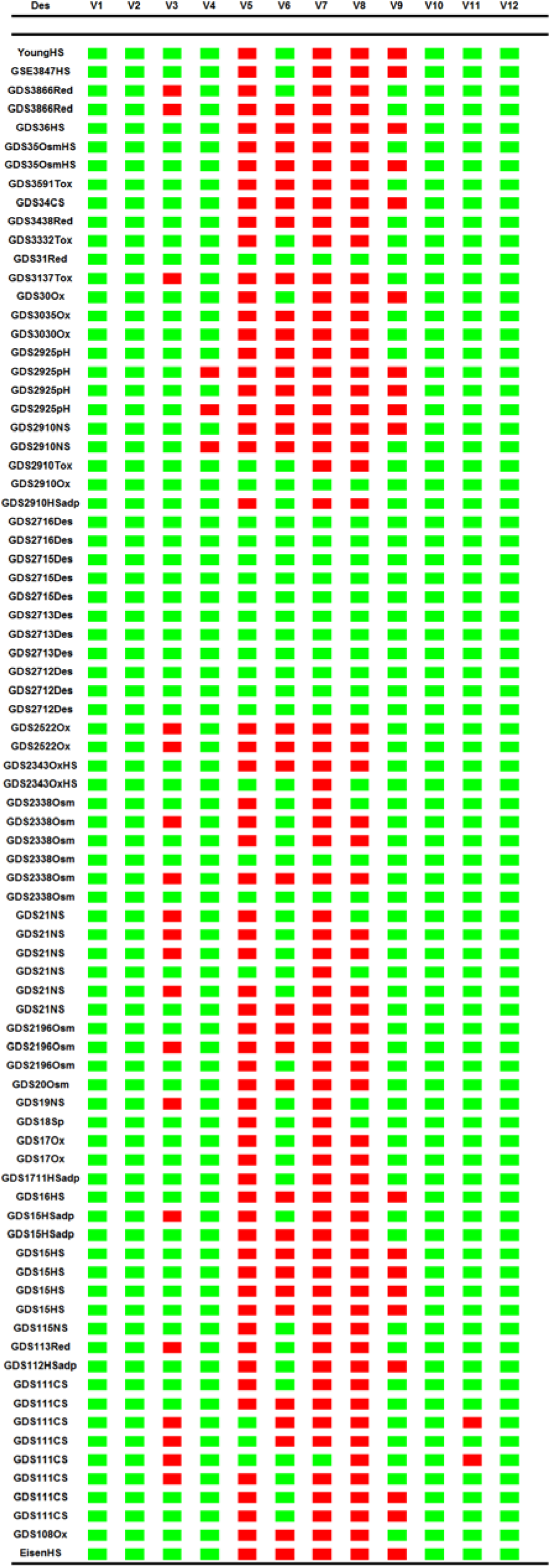
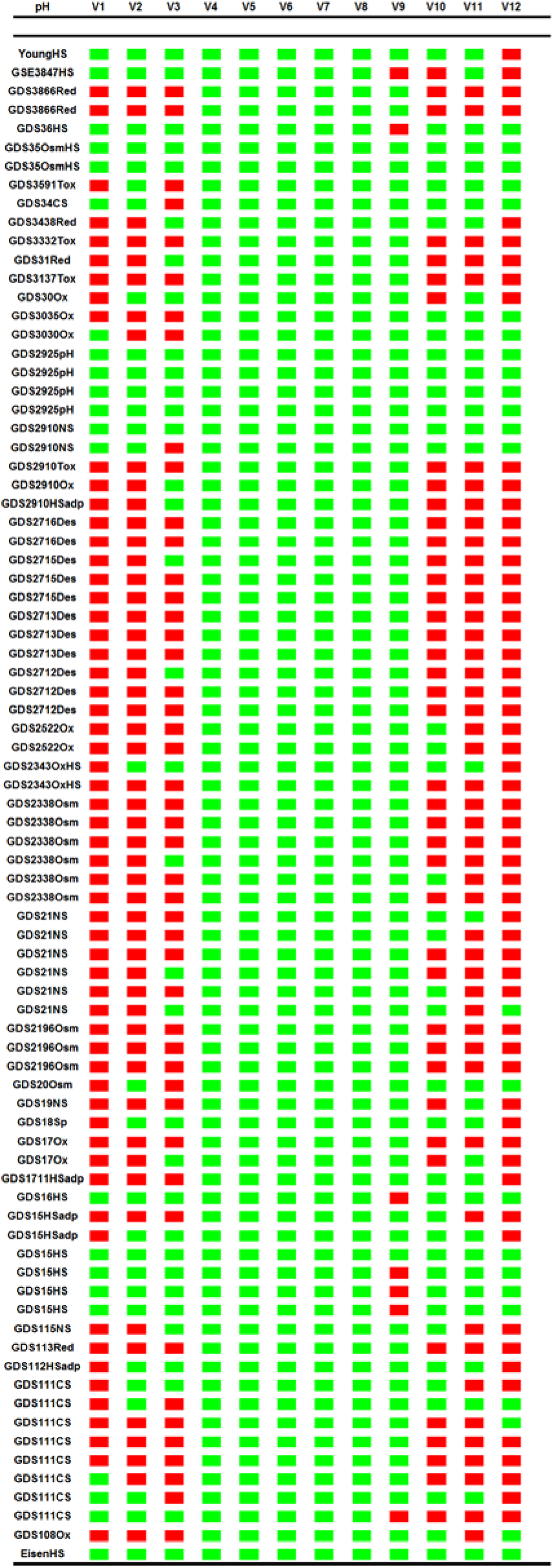
Heat map representation of the feasibility space for physiological variables V1-V11. Each column represents a variable from Table I of the main manuscript, while each row represents a database of gene expression. A – Feasibility space for desiccation-rehydration response. The eleven variables defined in Table I of the main manuscript were calculated for the data bases that measured changes in gene expression during response to Desiccation-Rehydration. The maximum and minimum values for each variable were recorded and used to build a feasibility space as described in the main manuscript. For each database, if the variable falls within (outside) this range, we color the entry of the heat map as green (red). A stress response falls within the desiccation/rehydration feasibility space if all entries in a row are green. We see that, with one exception (response to strong osmotic stress) only desiccation-rehydration shock responses fall fully within this feasibility space. B – Feasibility space for pH stress response. The eleven variables defined in Table I of the main manuscript were calculated for the data bases that measured changes in gene expression during response to pH shift. The maximum and minimum values for each variable were recorded and used to build a feasibility space as described in the main manuscript. For each database, if the variable falls within (outside) this range, we color the entry of the heat map as green (red). A stress response falls within the pH feasibility space if all entries in a row are green. We see that mostly pH responses fall fully within this feasibility space.

**Supplementary Figure 3.**
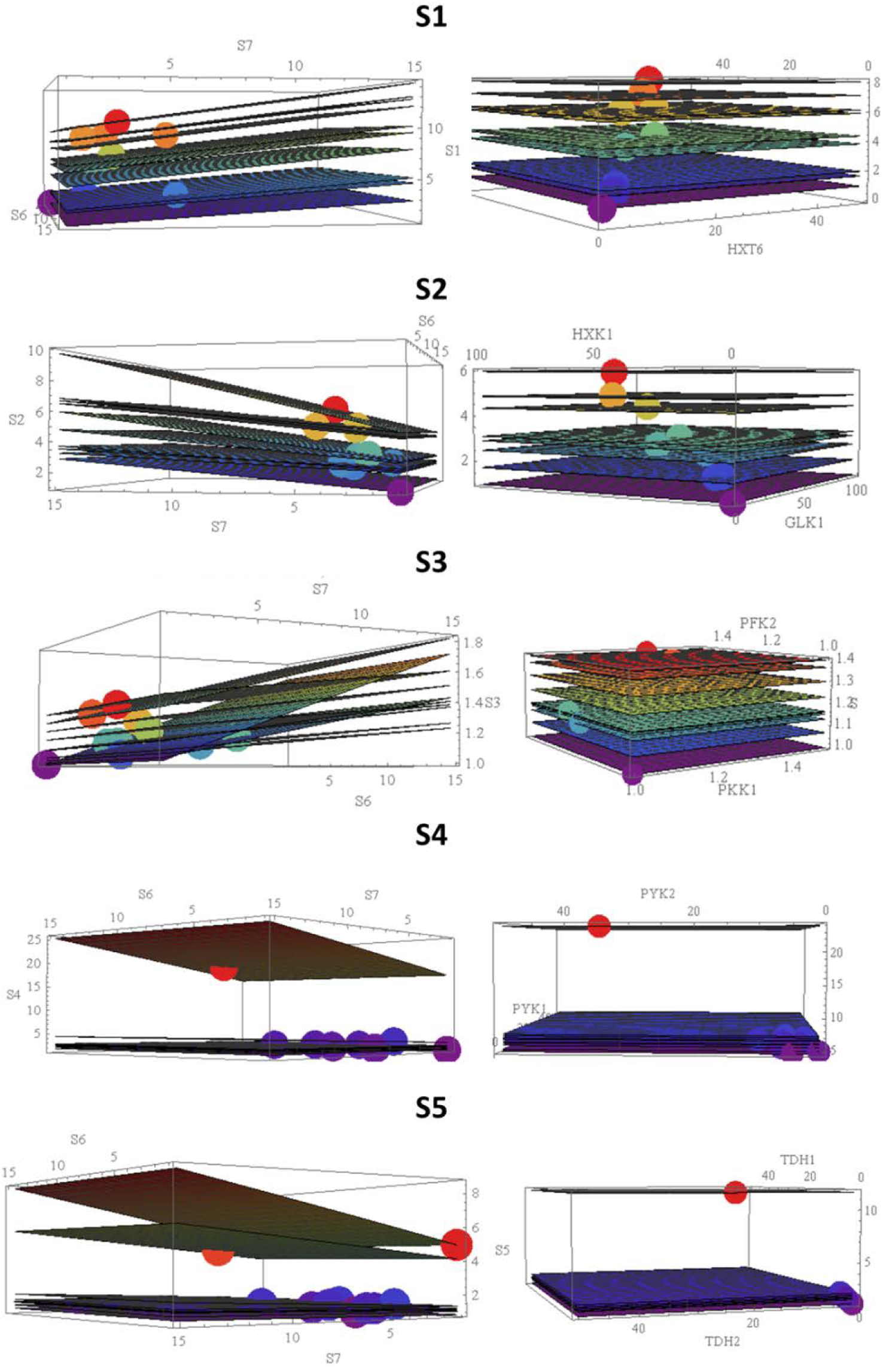
Evolution can find different combinations of changes in gene expression and enzyme activities that are equivalent with respect to the changes they cause in variables V1 – V1 1. Graphical representation of this situation for activities S1 – S5. Each plane represents one of the heat shock response databases used to calculate the feasibility region shown in Figure 1A. Left Column Activities S1-S5 as a function of activities S6 and S7. Each plane represents all possible sets of values for S1-S5, S6, and S7 that would generate the same values for V1 – V11 for the same heat shock database. The dot that falls in each plane represents the actual activity estimated from the experimental changes in gene expression data. Right Column – Activities S1-S5 as a function of genes that code for proteins that contribute for that activity, considering the same heat shock databases we use as an example in the left column. Each plane corresponds to one of the databases. All points falling on a plane are formally equivalent, leading to the same activity. The dots in each plane represent the actual measurement for the adaptive response.

**Supplementary Figure 4.**
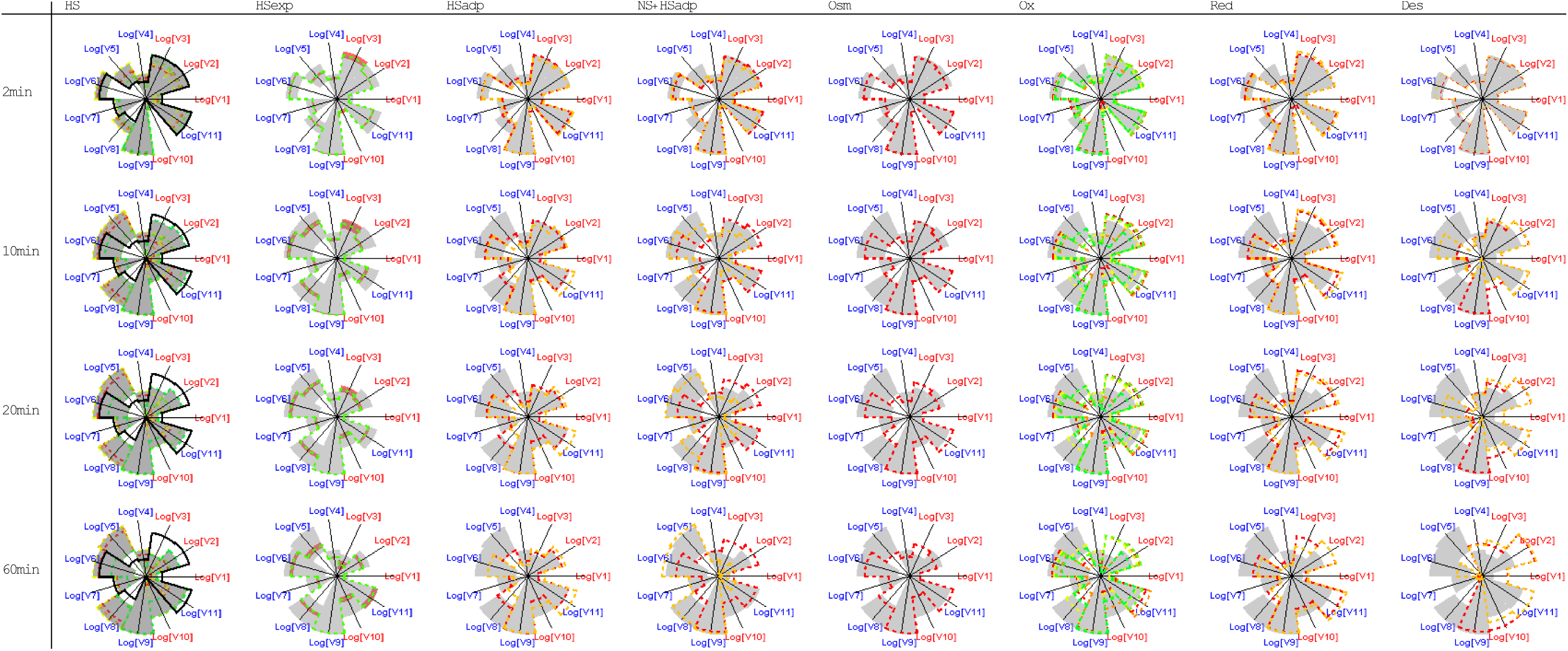
Spider plot representation for the dynamic adaptation of the eleven physiological variables from Table I during the time course of heat shock response. The time course was generated as described in methods and using the databases marked with “*” in Supplementary Table II. We show snapshots of the adaptation at 2, 10, 20 and 60 min after the onset of the environmental challenge. The grey areas are defined by the heat shock responses at each time (HS column). The “HSExp” column shows the median results for our time course measurements of heat shock adaptation, also representing quantiles 0.25 to 0.75 around that median value. The “NS+HS” column shows the time course adaptation for cells preadapted to nutrient stress followed by heat shock adaptation. The “Osm” column shows the time course adaptation for osmotic shock. The “Ox” column shows the time course adaptation for oxidative stress. The “Red” column shows the time course adaptation for reductive stress. The “Des” column shows the time course adaptation for desiccation/rehydration stress.

## Supplementary online material for Metabolic constraints and quantitative design principles in gene expression during adaption of yeast to heat shock

### 1 Supplementary Methods

#### 1.1 Biological System to Model

From years of accumulated research, it is apparent that in order to adapt to heat shock, *Saccharomyces cerevisiae* has to adjust its production of different metabolic variables. ^1,2^. These adjustments are described in Table 1 of the main manuscript. Considering that exponentially growing *S. cerevisiae* in a rich medium use glycolysis as the main energy pathway, we have defined a minimal representation of the molecular system that needs to be modelled in order to estimate the changes in all the physiological variables of Table 1 as the cell adapts to stress. This representation is shown in Supplementary Figure 1.

Experimentally derived parameter values for this model were taken from ^1,2^ and are given in Eqs 1-5 of the main manuscript. The basal model is provided as an SBML file and in supplementary notebooks 1-6. It can be used as described in the main methods section to calculate the effect of changes in gene expression on the variables described in Table 1 of the main manuscript. All computations were done using Mathematica™ ^3^ and the open-source package MathSBML ^4^.

#### 1.2 Modeling Formalism

In 1969 Michael Savageau proposed an approximate formalism that can be used to build mathematical models of systems for which only limited kinetic information is available ^5,6^. This formalism uses the Taylor theorem, which states that any continuously differentiable function can be exactly represented by a polynomial series of its variables. If one creates the Taylor series in a non-linear logarithmic space and truncates the series at the first order term, upon return to Cartesian space, one obtains a power law representation of the function.

Assume that the mathematical model that describes the dynamical behavior of a biological system can be written as the following set of ordinary differential equations (ODEs):

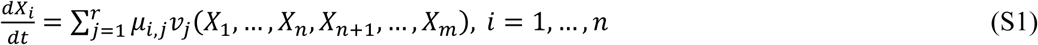

In these ODEs *v_j_* represents all the rates of the individual processes in the system and *μ_i,j_* is the stoichiometric coefficient of *X_i_* in process of reaction *j*. It is positive if flux *j* produces *X_i_*, negative if flux *j* depletes the pool of *X_i_*, and zero if *X_i_* is neither produced nor consumed by flux *j*. *X*_1_,…, *X_n_* are the dependent variables of the system and *X_n_*_+1_,…, *X_m_* are the independent variables of the system.

One can use a Generalized Mass Action (GMA) representation within the power law formalism described above ^1,7,8^, and write the differential equations for each of the n dependent variables of the system as:

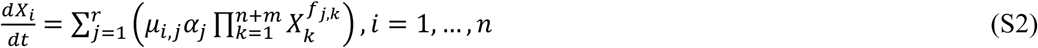

where *X_i_* represents the concentration of metabolite *i* in the model, *m* is the number of independent variables, and *r* is the number of fluxes in the system. *α_j_* is the apparent rate constant of reaction *j*. *f_j,k_* is the kinetic order of variable *X_k_* in reaction *j*. Each kinetic order quantifies the effect of the metabolite *X_k_* on flux *j* and corresponds to the local sensitivity of the rate *v_j_* to *X_k_*, evaluated at the operating point indicated by the subscript 0:

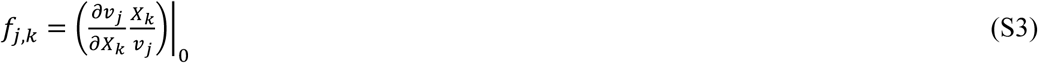

If *X_k_* has no direct influence on the rate of reaction *j*, the kinetic order is zero. If *X_k_* directly activates the flux of reaction *j*, the kinetic order is positive. If *X_k_* directly inhibits the flux of reaction *j*, the kinetic order is negative.

#### 1.3 Sensitivity Analysis

Biological systems must be able to adapt to and survive in an ever-changing environment, while maintaining low sensitivity to spurious fluctuation in parameters (e.g. enzyme activity or *Km*) that should not greatly affect its steady state or homeostasis ^5,9,10^. This is called robustness of the steady state and it can be measured using sensitivity analysis. In this work we evaluated the robustness of the physiological variables V1-V11 using relative steady state parameter sensitivities. Relative steady state parameter sensitivities are “the relative change in a system component (X) that is caused by a relative change in a parameter value (p),” ^11^:

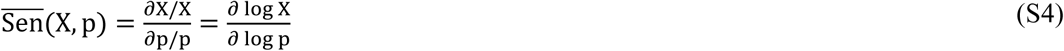

The extension of this definition for a system of ODEs is trivial and can be seen in ^11^. In our case, we wanted to determine the relative sensitivity of the physiological variables V_1_ – V_11_ of Table I to the enzyme activities S1 – S7:

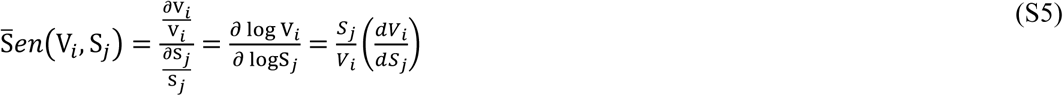

The results of this sensitivity analysis identify the enzyme activities that more strongly influence each of the relevant physiological variables. In approximate terms, if 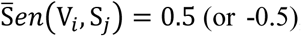, this means that when the value of S_j_ changes by 100%, the value of V_i_ is expected to increase (or decrease) by 50%. Supplementary notebook I provides the script for all these calculations.

#### 1.4 Steady state stability

The stability of the steady states was determined using linear stability analysis. First we calculated the Jacobian matrix for the differential equations (Eqs 1-5 of the main manuscript) which is defined as

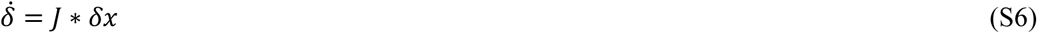

Where J* is the Jacobian evaluated at the equilibrium point, the solution is the set of eigenvalues of the Jacobian. Then, for each steady state, we calculated the five eigenvalues of that matrix. If the steady state is stable, all eigenvalues have negative real parts ^5,9,10^. Supplementary notebook I provides the script for all these calculations.

### 2 Supplementary Results

#### 2.1 Sensitivity and Stability Analysis

Sensitivity analysis estimates how dependent variables and outputs of the model change with respect to changes and/or fluctuations in the values of parameters and independent variables of the model. High sensitivities tend to correlate well with parts of the model that are less accurate representations of real phenomena. We used relative sensitivities (see main methods section) to analyze the impact of changes in enzyme activities S1 - S7 of Supplementary Table IV in the eleven physiological variables of interest described in Table I of the main manuscript.

We calculated relative sensitivities of the steady state for all microarray experiments. Results are summarized in Supplementary Table VI, below. We now briefly describe the activities that have the stronger influence in each physiological variable V1-V11.

V1 (VATP) is positively affected by increases in the activity of S1. V2 (VTre) is positively affected by increases in activities S1 and S6. It is also inversely affected by increases in S3. In addition, increases in S7 were also predicted to cause significant decreases in V2 in some cases (mostly in response to desiccation). V3 (VNADPH) is positively affected by increases in S7. V4 (Glucose levels) is positively affected by increases in S1 and negatively affected by increases in S2. In a small number of cases V4 is also positively influenced by increases in S7. V5 (Glucose-6-phosphate levels) is positively affected by increases in S1 and negatively influenced by increases in S3. In a small number of cases V5 is also negatively influenced by increases in S7. V6 (Fructose 1-6 Bisphosphate) is positively affected by increases in S1 and negatively affected by increases in S4. V7 (Phosphoenolpyruvate) is positively affected by increases in S1 and negatively affected by increases in S5. In addition, in desiccation experiments, V7 is positively affected by increases in S3 and negatively affected by increases in S7. V8 (ATP) is positively affected by increases in S1. Increases in S7 (S3) negatively (positively) affect V8 in some adaptive response, mostly to desiccation. V9 (a proxy for biosynthetic cost of the response) does not show an overall pattern.

However, if it is strongly affected by one the activities, it is also strongly affected by all the others. V10 (VGLY) is positively affected by S1 and negatively affected by S5. V11 (ψ) is positively affected by S3 (and in some cases by S7), and negatively affected by the S1.

Stability analysis of the steady state is a valuable predictor of the qualitative behavior of the system. Systems with stable steady states are more likely to be adequate representation of biological phenomena. At steady state, the eigenvalues of the Jacobian matrix of the ODE system that represent the biological process under study measure this stability. Eigenvalues are typically complex numbers with an imaginary and real part. If all the eigenvalues have negative real parts, the systems will have a locally stable steady-state ^10^. The steady states for all microarray experiments are locally stable, as they all have only eigenvalues with negative real parts.

All calculations can be reproduced using Supplementary Notebook 1.

#### 2.2 Mapping phenotype to genotype

Our mathematical model can be used to calculate five enzyme activities as functions of the dependent variables and of the remaining two enzyme activities. This is shown in Eqs. S4 –S8, where S1-S5 are calculated as a function of S6 and S7:

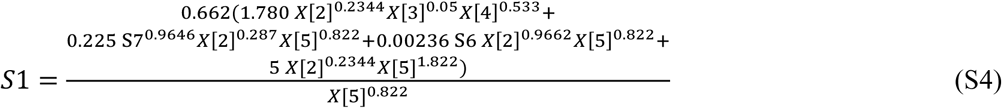

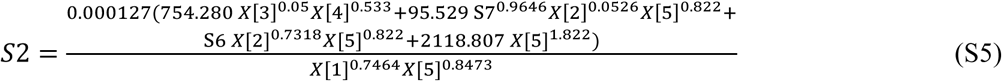

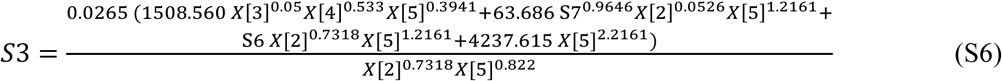

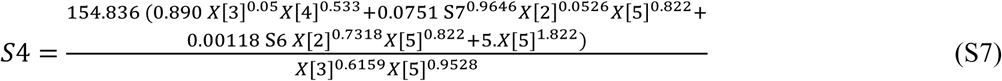

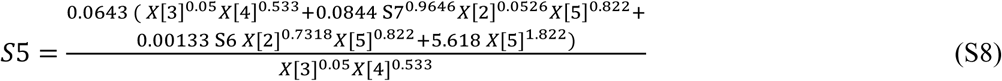

#### 2.3 Correlation between changes in gene expression and changes in protein abundance during heat shock response

In order to evaluate how accurate our assumption of direct proportionality between changes in gene expression and changes in protein levels is for our genes of interest we downloaded the whole proteome abundance measurements made by Ghaemmaghami *et al.*^12^. Then, we downloaded the data from CYCLoPS ^13,14^, and from Mackenzie et al. ^15^ pertaining to the whole proteome quantification in *S. cerevisiae* after heat shock. Finally, we calculated the median of the ratios between protein abundance after heat shock with respect to protein abundance under basal conditions. In parallel, we calculated the median change in gene expression from each gene of interest to the model in the heat shock databases. The results are shown here:

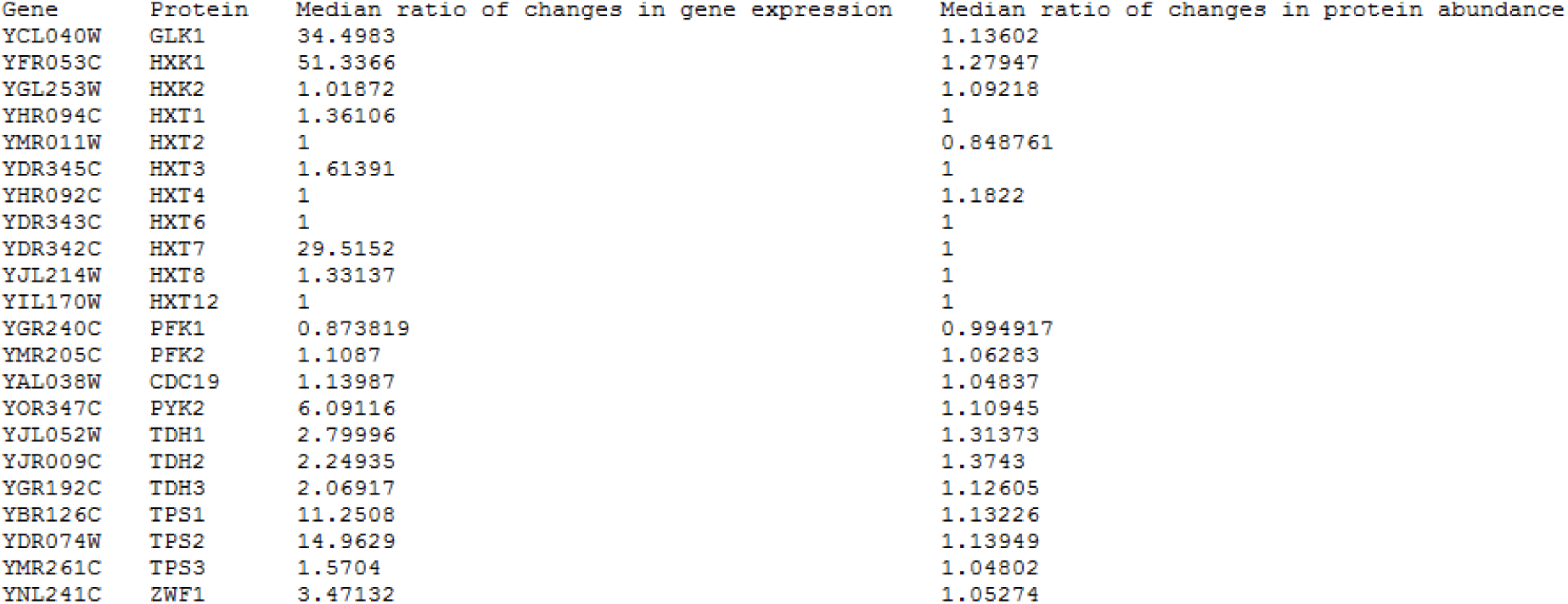

| Gene | Protein | Median ratio of changes in gene expression | Median ratio of changes in protein abundance |
| --- | --- | --- | --- |
| YCL040W | GLK1 | 34.4983 | 1.13602 |
| YFR053C | HXK1 | 51.3366 | 1.27947 |
| YGL253W | HXK2 | 1.01872 | 1.09218 |
| YHR094C | HXT1 | 1.36106 | 1 |
| YMR011W | HXT2 | 1 | 0.848761 |
| YDR345C | HXT3 | 1.61391 | 1 |
| YHR092C | HXT4 | 1 | 1.1822 |
| YDR343C | HXT6 | 1 | 1 |
| YDR342C | HXT7 | 29.5152 | 1 |
| YJL214W | HXT8 | 1.33137 | 1 |
| YIL170W | HXT12 | 1 | 1 |
| YGR240C | PFK1 | 0.873819 | 0.994917 |
| YMR205C | PFK2 | 1.1087 | 1.06283 |
| YAL038W | CDC19 | 1.13987 | 1.04837 |
| YOR347C | PYK2 | 6.09116 | 1.10945 |
| YJL052W | TDH1 | 2.79996 | 1.31373 |
| YJR009C | TDH2 | 2.24935 | 1.3743 |
| YGR192C | TDH3 | 2.06917 | 1.12605 |
| YBR126C | TPS1 | 11.2508 | 1.13226 |
| YDR074W | TPS2 | 14.9629 | 1.13949 |
| YMR261C | TPS3 | 1.5704 | 1.04802 |
| YNL241C | ZWF1 | 3.47132 | 1.05274 |

We then calculated the Spearman’s Rho between the ratio of changes in protein abundance and gene expression. We found that this correlation coefficient was 0.55, which indicate a very strong correlation between changes in gene expression and changes in protein abundance.

---

**Supplementary Table I.**
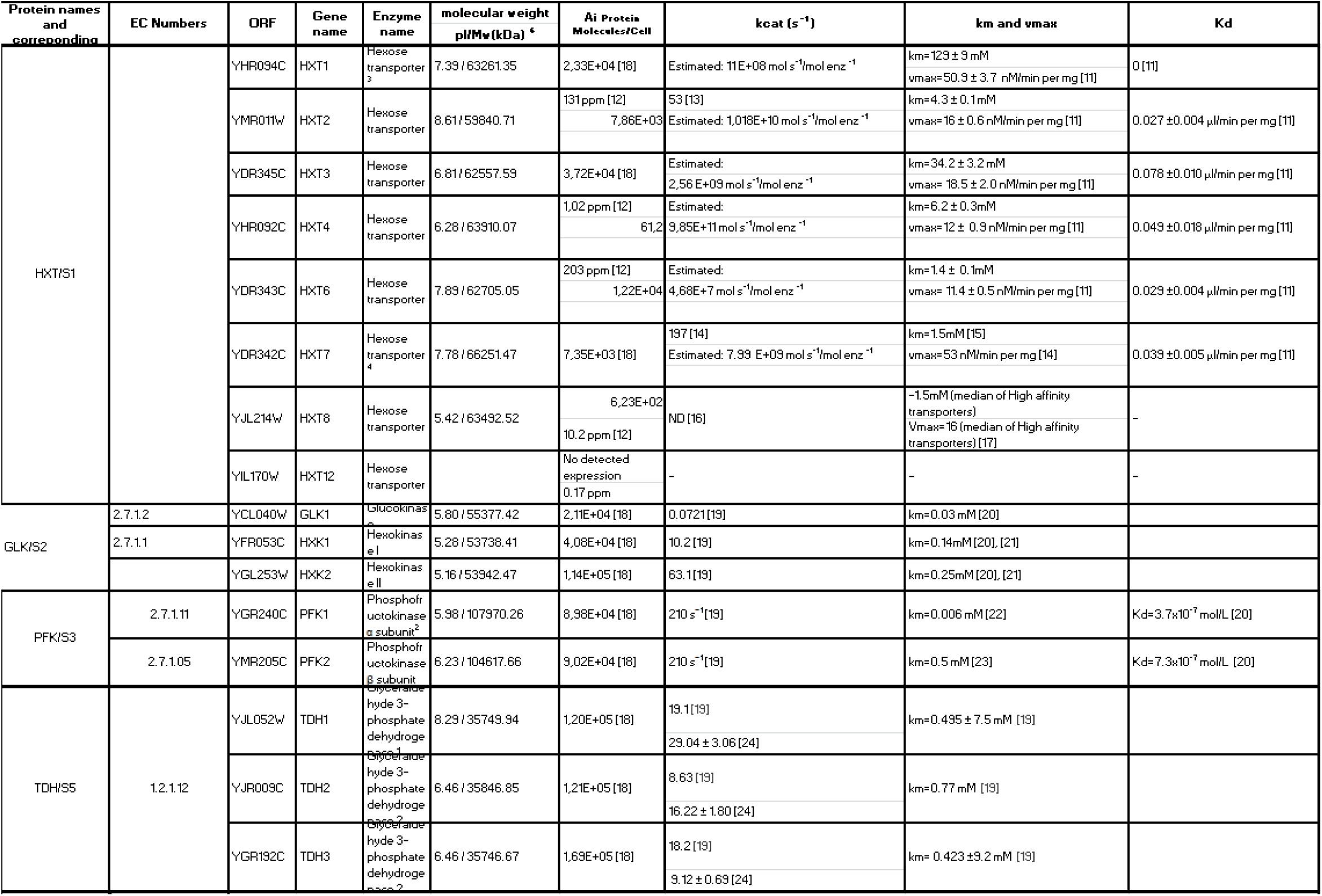
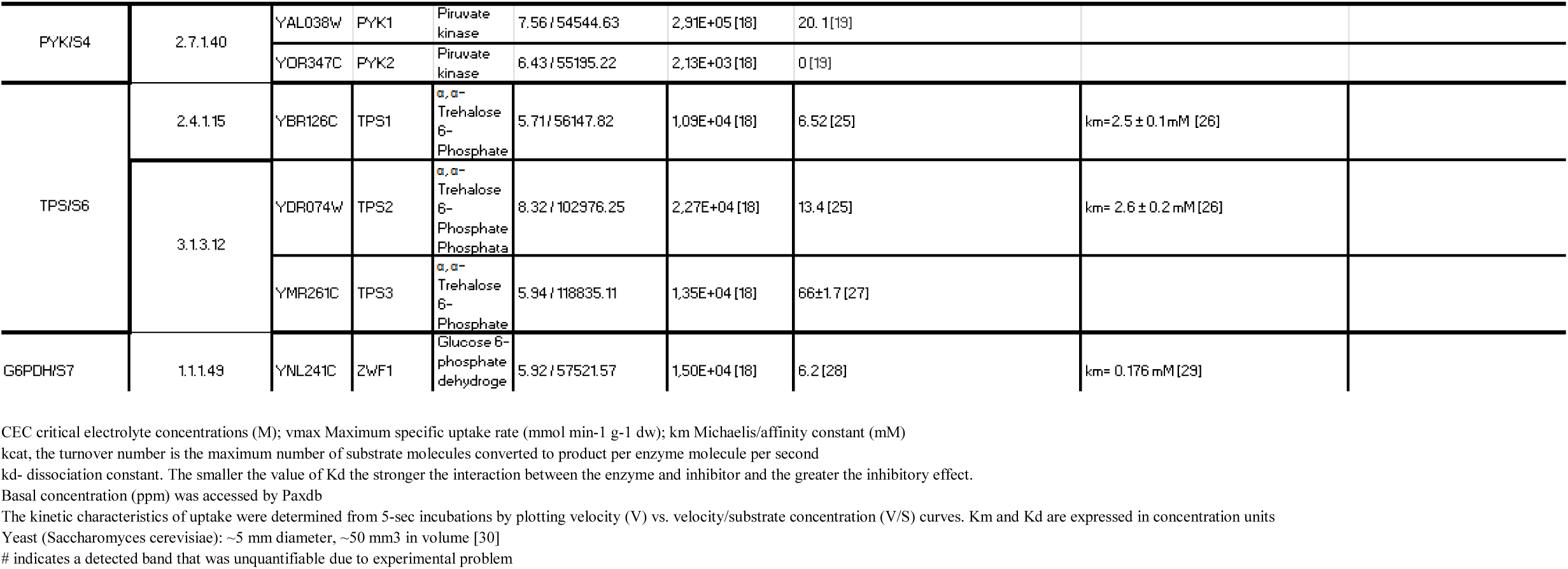
Enzyme activities, EC numbers and genes coding for enzymes relevant to the mathematical model described by Eqs 6-10 of the main manuscript.

**Supplementary Table II.**
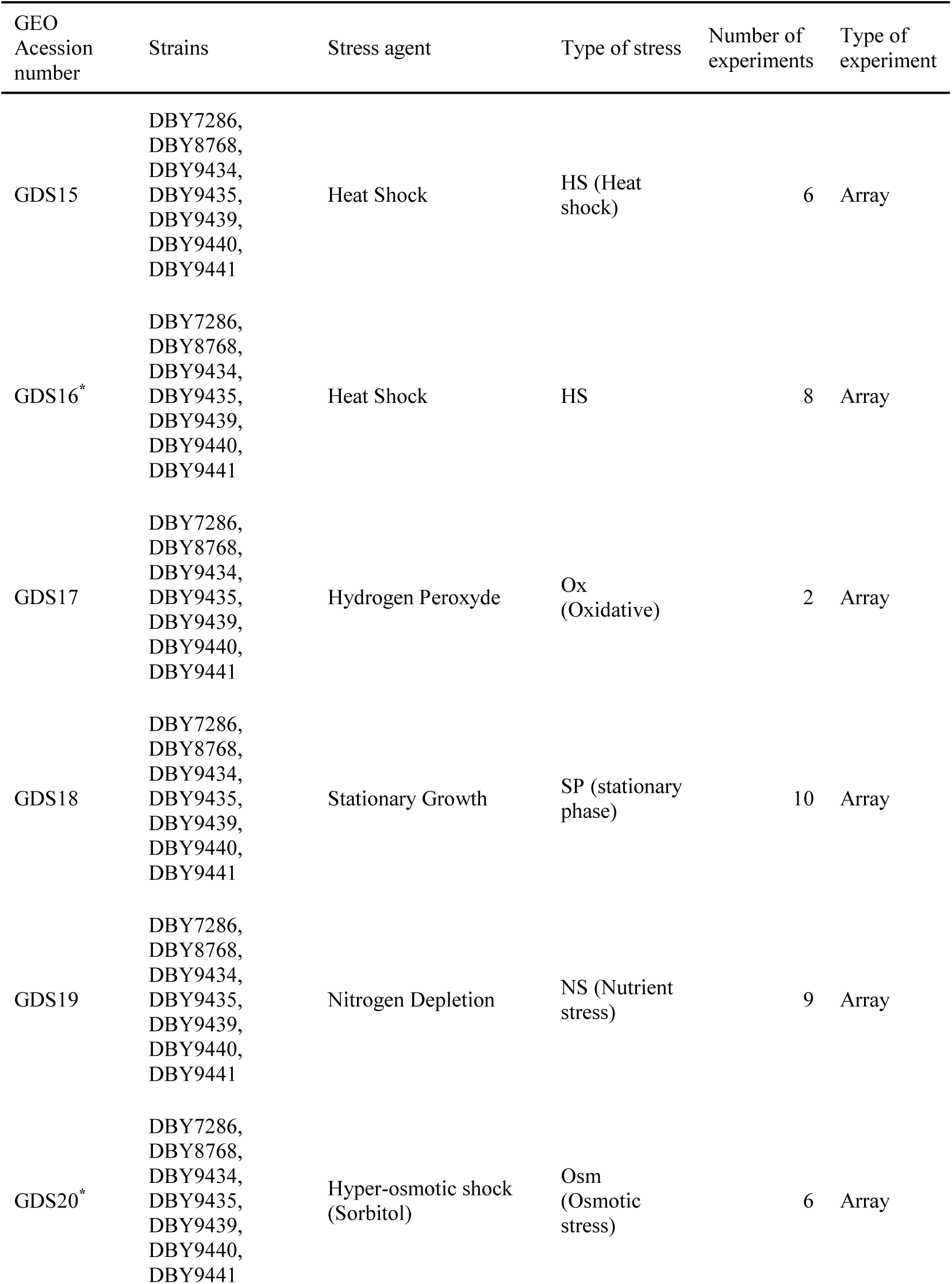
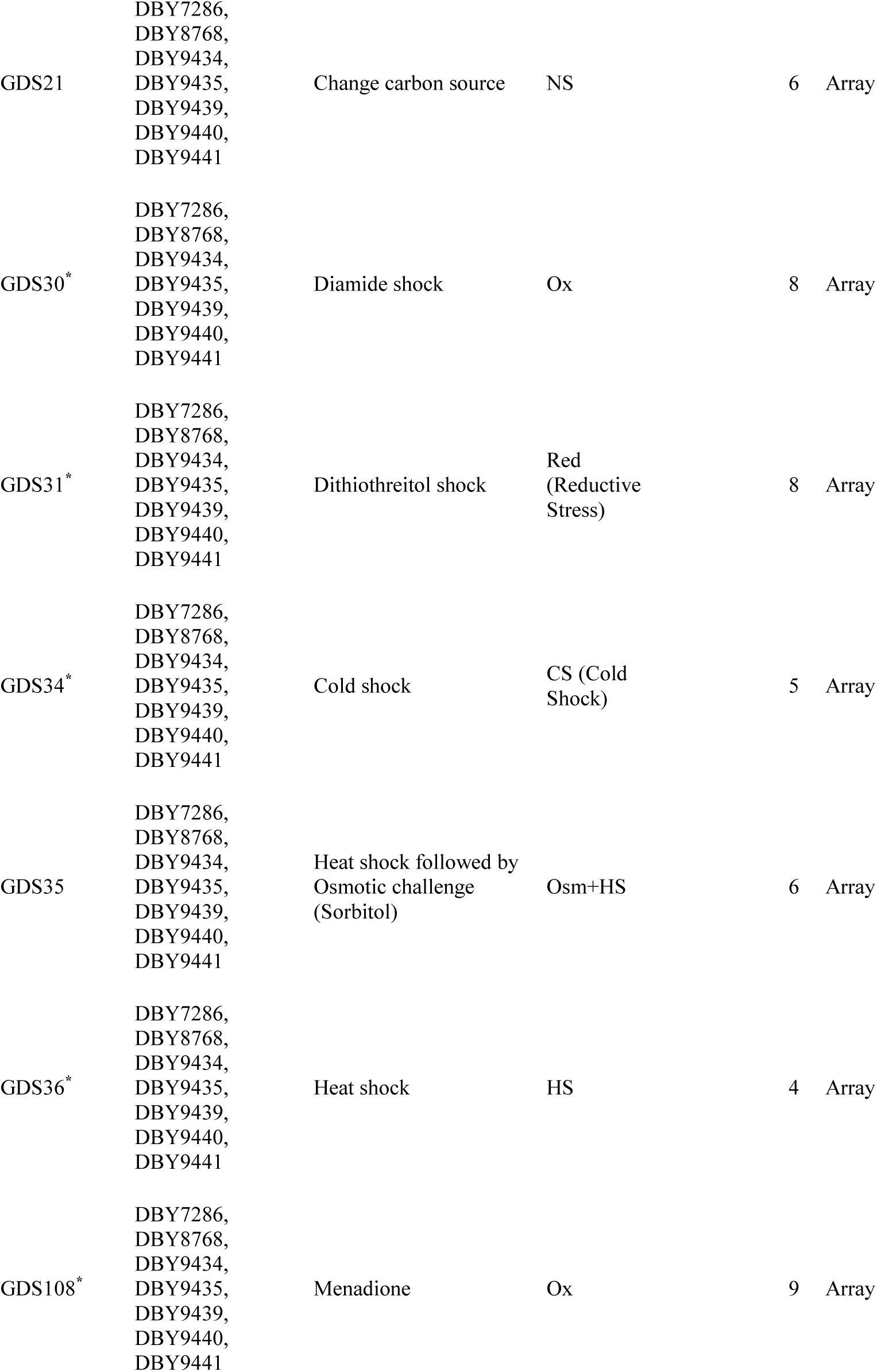
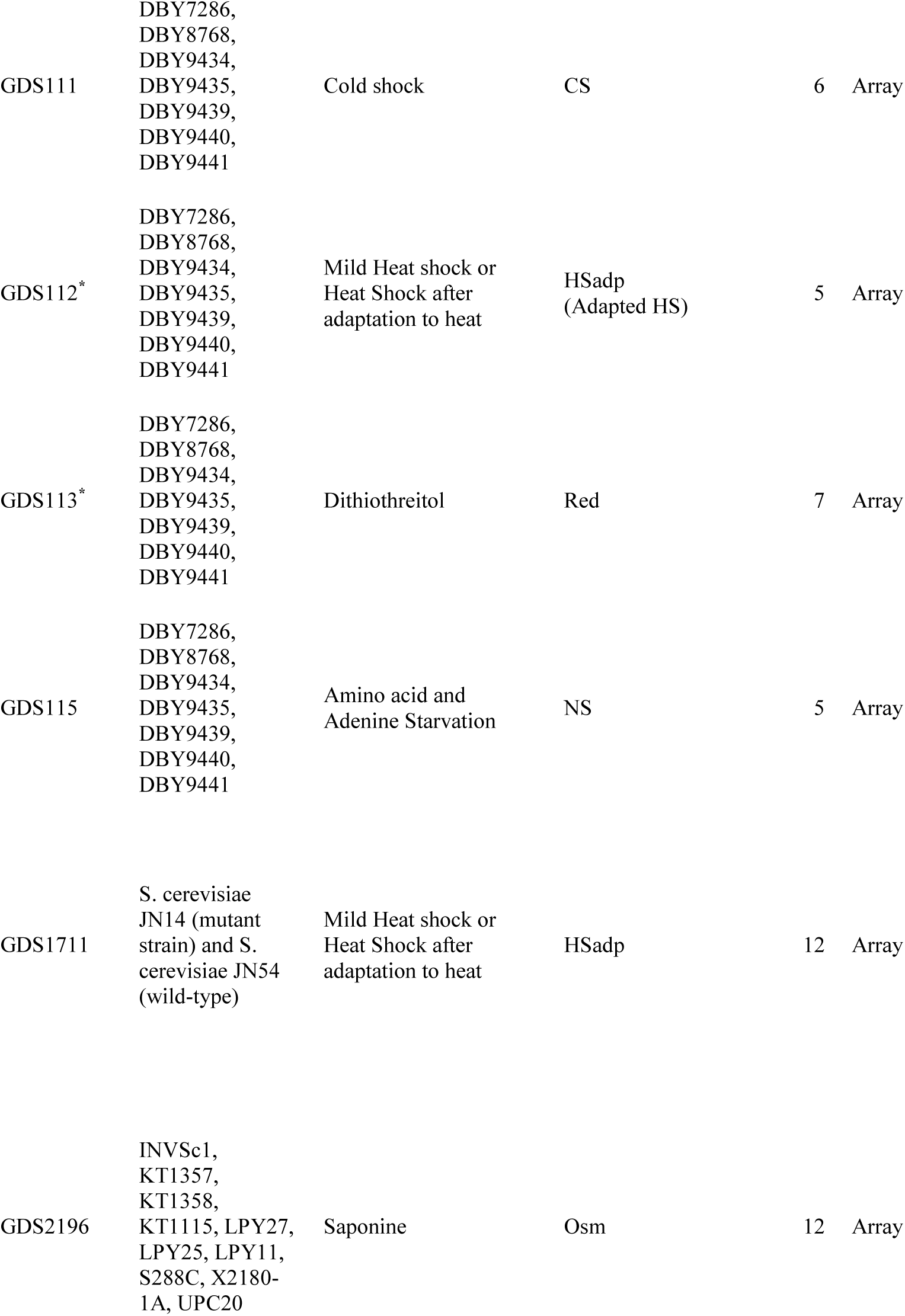
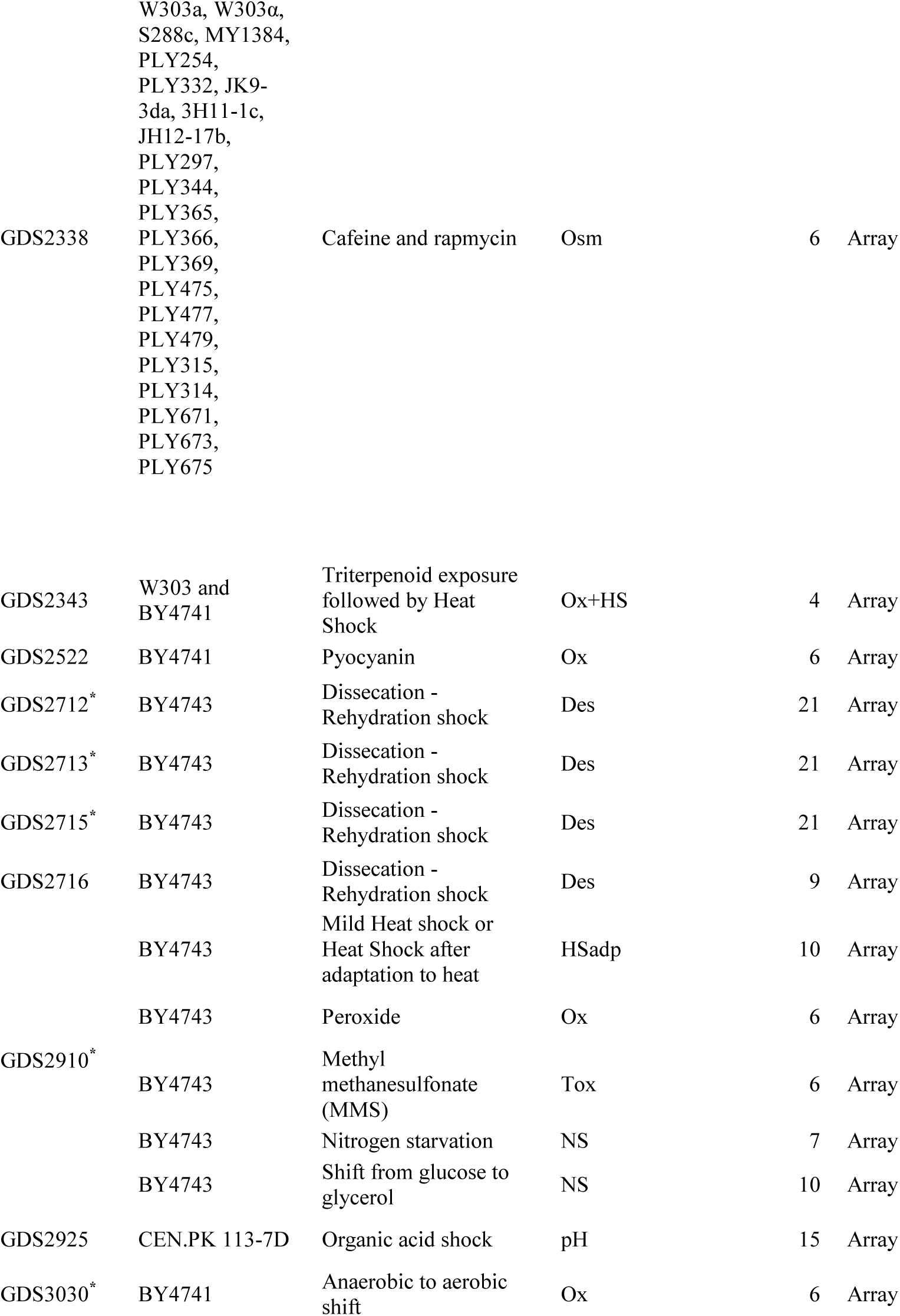
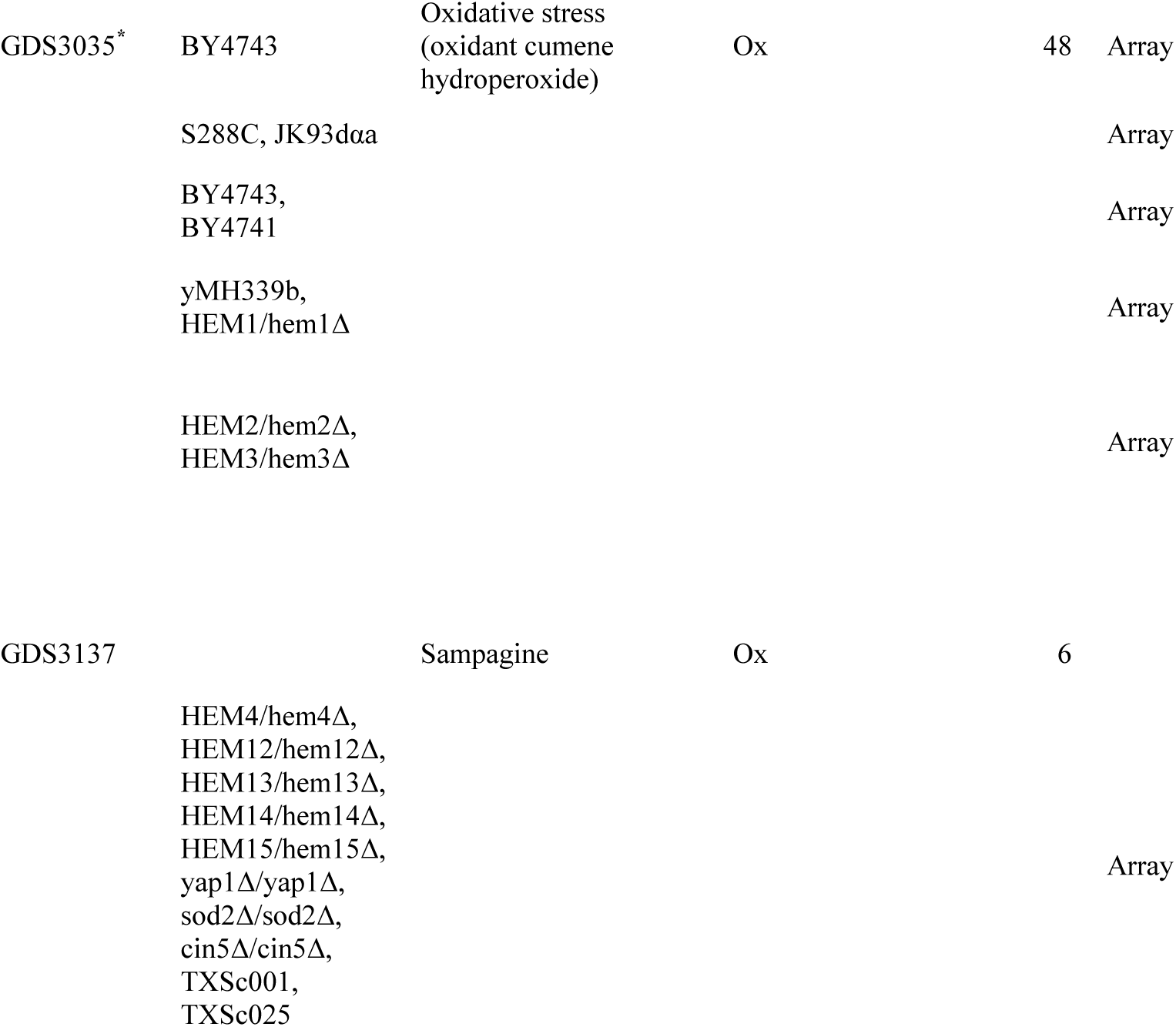
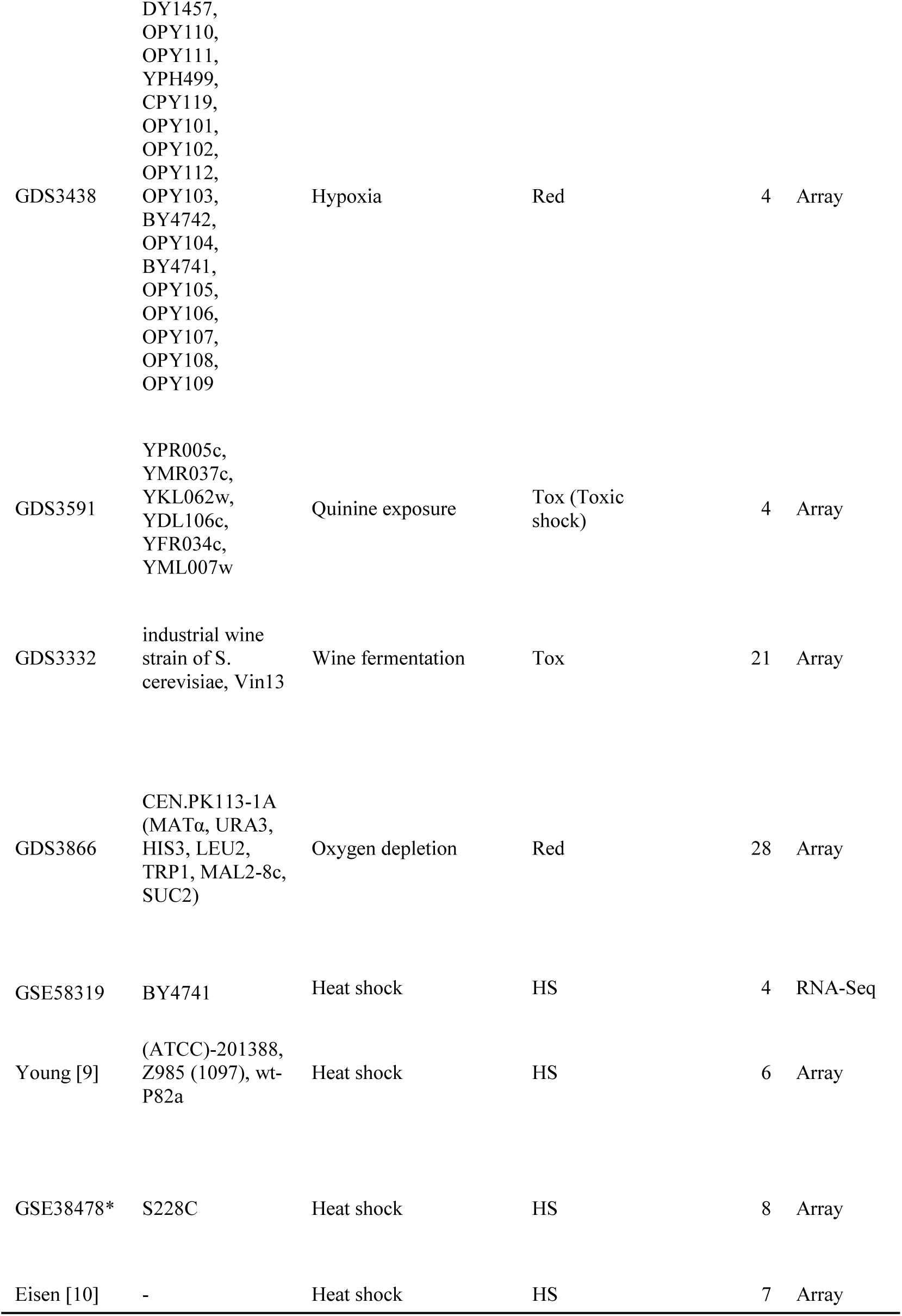
*S. cerevisiae* gene expression datasets downloaded from GEO or the primary literature and used in our analysis. Databases marked with “*” contained time course information that was used to generate Supplementary Figure 4 and Table IV.

**Supplementary Table III.**
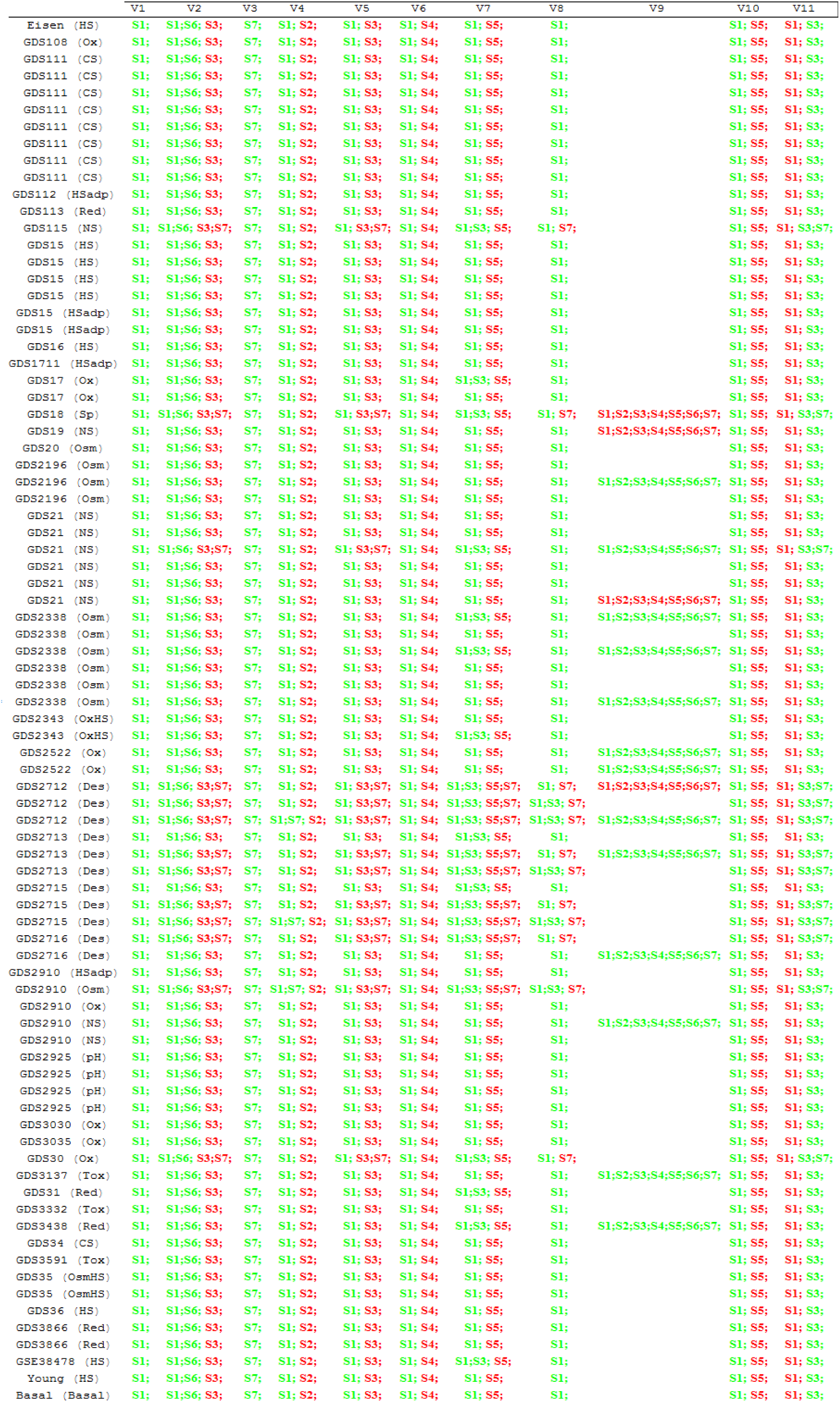
Relative Sensitivities of physiological variables V1-V11 to enzyme activities S1-S7. Red indicates sensitivities lower than -0.5. Green indicates sensitivities higher than 0.5. Sensitivities between -0.5 and 0.5 are not represented.

**Supplementary Table IV.**
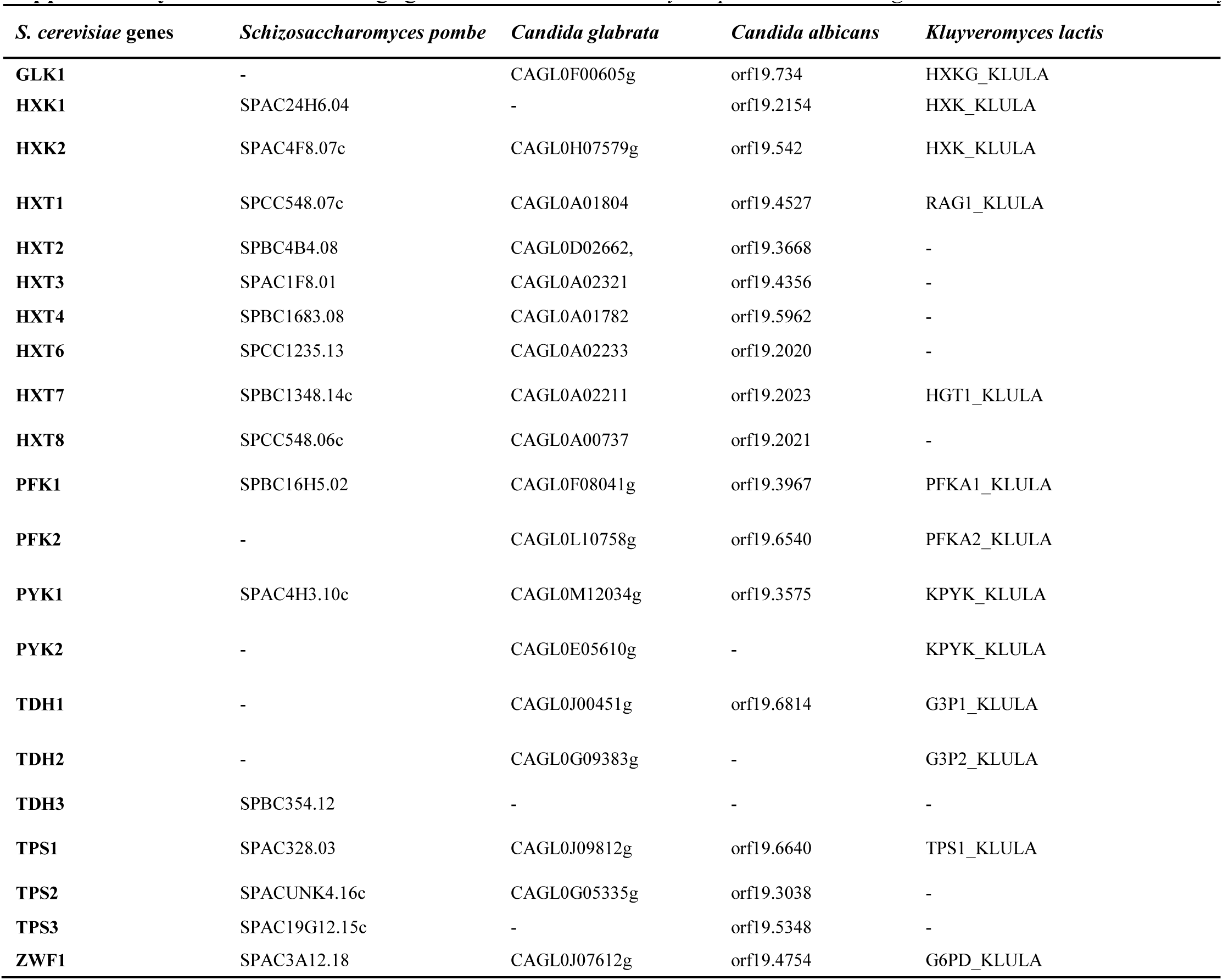
Orthologs genes in *Schizosaccharomyces pombe, Candida glabrata, Candida albicans, Kluyveromyces lactis*.

| *S. cerevisiae* genes | *Schizosaccharomyces pombe* | *Candida glabrata* | *Candida albicans* | *Kluyveromyces lactis* |
| --- | --- | --- | --- | --- |
| **GLK1** | - | CAGL0F00605g | orf19.734 | HXKG_KLULA |
| **HXK1** | SPAC24H6.04 | - | orf19.2154 | HXK_KLULA |
| **HXK2** | SPAC4F8.07c | CAGL0H07579g | orf19.542 | HXK_KLULA |
| **HXT1** | SPCC548.07c | CAGL0A01804 | orf19.4527 | RAG1_KLULA |
| **HXT2** | SPBC4B4.08 | CAGL0D02662, | orf19.3668 | - |
| **HXT3** | SPAC1F8.01 | CAGL0A02321 | orf19.4356 | - |
| **HXT4** | SPBC1683.08 | CAGL0A01782 | orf19.5962 | - |
| **HXT6** | SPCC1235.13 | CAGL0A02233 | orf19.2020 | - |
| **HXT7** | SPBC1348.14c | CAGL0A02211 | orf19.2023 | HGT1_KLULA |
| **HXT8** | SPCC548.06c | CAGL0A00737 | orf19.2021 | - |
| **PFK1** | SPBC16H5.02 | CAGL0F08041g | orf19.3967 | PFKA1_KLULA |
| **PFK2** | - | CAGL0L10758g | orf19.6540 | PFKA2_KLULA |
| **PYK1** | SPAC4H3.10c | CAGL0M12034g | orf19.3575 | KPYK_KLULA |
| **PYK2** | - | CAGL0E05610g | - | KPYK_KLULA |
| **TDH1** | - | CAGL0J00451g | orf19.6814 | G3P1_KLULA |
| **TDH2** | - | CAGL0G09383g | - | G3P2_KLULA |
| **TDH3** | SPBC354.12 | - | - | - |
| **TPS1** | SPAC328.03 | CAGL0J09812g | orf19.6640 | TPS1_KLULA |
| **TPS2** | SPACUNK4.16c | CAGL0G05335g | orf19.3038 | - |
| **TPS3** | SPAC19G12.15c | - | orf19.5348 | - |
| **ZWF1** | SPAC3A12.18 | CAGL0J07612g | orf19.4754 | G6PD_KLULA |

**Supplementary Table V.** GEO databases measuring gene expression changes during heat shock response for *S. pombe, K. lactis, C. albicans*, and *C. glabrata*.

| DB name | Specie | Strain | Condition of stress | Type of stress | Number of experiments |
| --- | --- | --- | --- | --- | --- |
| GSE50156 | Schizosaccharomyces pombe | SO3483,<br>SO4281 | Heat Shock | HS | 1 |
|  |  | SO4362,<br>SO4389 |  |  |  |
|  |  | SO4400,<br>SO5216 |  |  |  |
|  |  | SO5217,<br>SO5274 |  |  |  |
|  |  | SO5440,<br>SO5442 |  |  |  |
|  |  | SO5445,<br>SO5448 |  |  |  |
|  |  | SO5522,<br>SO5524 |  |  |  |
|  |  | SO5527,<br>SO5666 |  |  |  |
|  |  | SO5672,<br>SO5673 |  |  |  |
|  |  | SO5683,<br>SO5778 |  |  |  |
|  |  | SO5873,<br>SO5875 |  |  |  |
|  |  | SO5946,<br>SO5947 |  |  |  |
|  |  | SO6017,<br>SO6454 |  |  |  |
|  |  | SO6455,<br>SO6550 |  |  |  |
|  |  | SO6551,<br>SO6965 |  |  |  |
|  |  | SO7020,<br>SO7021 |  |  |  |
|  |  | SO7022,<br>SO7023 |  |  |  |
|  |  | SO7051,<br>SO7052 |  |  |  |
|  |  | SO7053,<br>SO7055 |  |  |  |
|  |  | SO7118 |  |  |  |
| GSE38478 | Schizosaccharomyces pombe |  | Heat Shock | HS | 8 |
| GSE38478 | Kluyveromyces lactis |  | Heat Shock | HS | 10 |
| GSE38478 | Candida glabrata |  | Heat Shock | HS | 11 |
| GSE38478 | Candida albicans |  | Heat Shock | HS | 10 |

